# *REST* and *RCOR* genes display distinct expression profiles in neurons and astrocytes using 2D and 3D human pluripotent stem cell models

**DOI:** 10.1101/2024.03.09.584254

**Authors:** Simon Maksour, Neville Ng, Amy J. Hulme, Sara Miellet, Martin Engel, Sonia Sanz Muñoz, Rachelle Balez, Ben Rollo, Rocio K. Finol-Urdaneta, Lezanne Ooi, Mirella Dottori

## Abstract

Repressor element-1 silencing transcription factor (REST) is a transcriptional repressor involved in neurodevelopment and neuroprotection. REST forms a complex with the REST corepressors, CoREST1, CoREST2, or CoREST3 (encoded by *RCOR1*, *RCOR2*, and *RCOR3*, respectively). Emerging evidence suggests that the CoREST family can target unique genes independently of REST, in various neural and glial cell types during different developmental stages. However, there is limited knowledge regarding the expression and function of the CoREST family in human neurodevelopment. To address this gap, we employed 2D and 3D human pluripotent stem cell (hPSC) models to investigate *REST* and *RCOR* gene expression levels. Our study revealed a significant increase in *RCOR3* expression in glutamatergic cortical and GABAergic ventral forebrain neurons, as well as mature functional NGN2-induced neurons. Additionally, a simplified astrocyte transdifferentiation protocol resulted in a significant decrease in *RCOR2* expression following differentiation. *REST* expression was notably reduced in mature neurons and cerebral organoids, along with *RCOR2* in the latter. In summary, our findings provide the first insights into the cell-type-specific expression patterns of *RCOR* genes in human neuronal and glial differentiation. Specifically, *RCOR3* expression increases in neurons, while *RCOR2* levels decrease in astrocytes. The dynamic expression patterns of *REST* and *RCOR* genes during hPSC neuronal and glial differentiation underscore the potential distinct roles played by REST and CoREST proteins in regulating the development of these cell types in humans.

**Graphical abstract:** 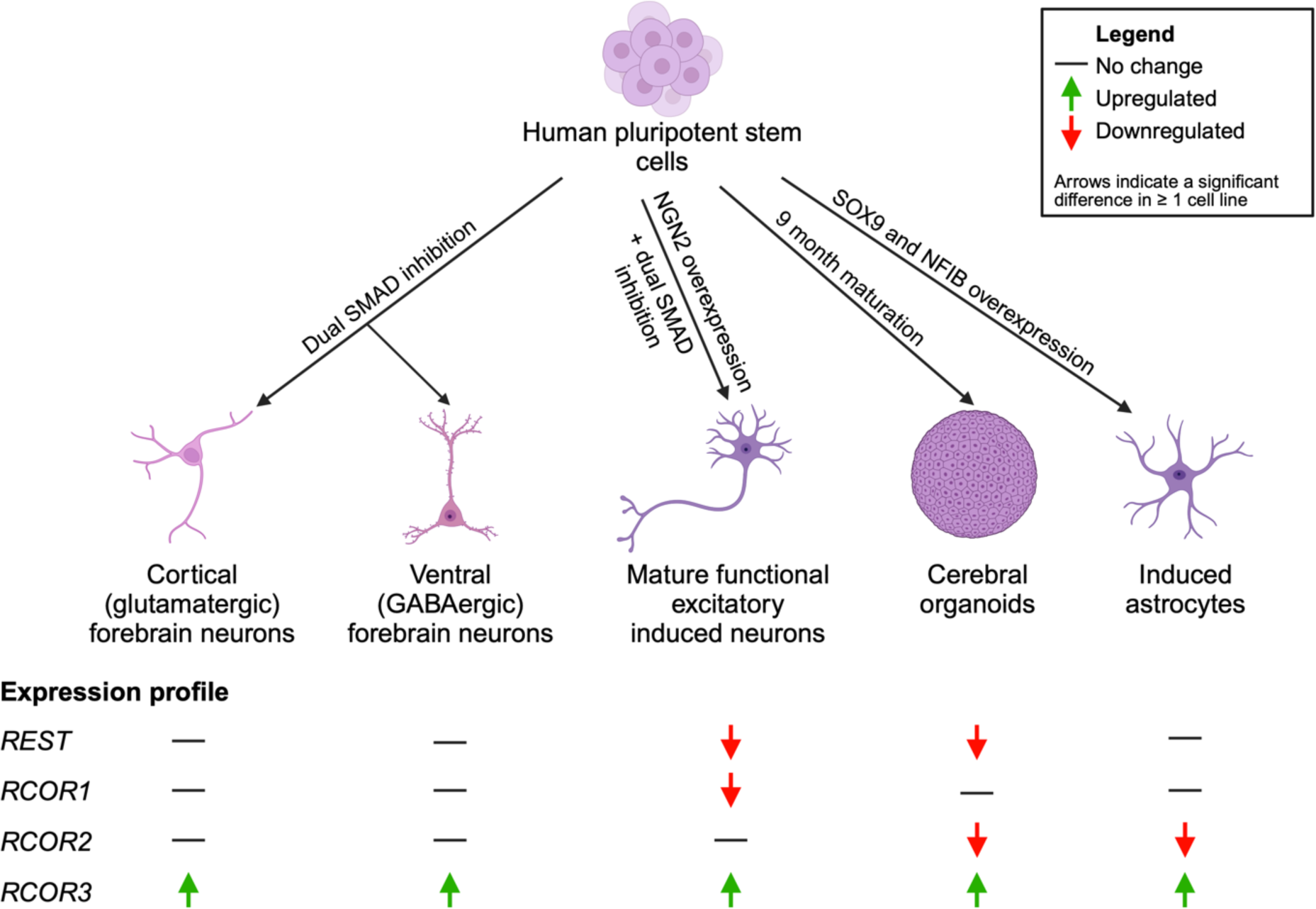

**Highlights:**

- *REST* and *RCOR* genes display cell-type specific expression patterns in neural cells
- *RCOR3* (encodes CoREST3) is upregulated during neuronal and astrocyte differentiation
- *RCOR2* (encodes CoREST2) is downregulated during differentiation of astrocytes
- Evidence of potential cell-type specific functions of the CoREST family

## Introduction

The molecular mechanisms orchestrating normal neurodevelopment are governed by timely and tightly regulated actions of transcription factor panels. Repressor element-1 silencing transcription factor (REST) is widely documented as a master regulator of neurogenesis, including roles in regulating neuronal subtype-specification and maturation [1,2], synaptic plasticity [3], somatosensory neuronal remodelling [4], and neuroprotection [5,6,7,8]. Genome wide analysis revealed approximately 2,000 potential target genes of REST in the human genome [9,10]. REST functions through the formation of a complex with REST corepressor 1 (CoREST1), CoREST2 or CoREST3 (encoded by *RCOR1*, *ROCR2* and *RCOR3*, respectively) and by recruiting chromatin modifying enzymes to induce a repressive chromatin state [11,12,13,14]. The importance of the CoREST protein family is just emerging as these proteins have the ability to bind to and repress the expression of unique target genes independently to REST, in various neural and glial cell types at different stages of development (as reviewed in Maksour, et al. (15)). Research has also begun to highlight the importance of the CoREST proteins as epigenetic regulators of neurogenesis, including a role in regulating pluripotency [14,16], neuronal differentiation and maturation [17,18,19], and neuroinflammation [20,21]. These findings indicate that each CoREST protein is more than simply a “REST corepressor” and has potentially distinct and critical roles in orchestrating the molecular mechanisms that govern neurodevelopment.

The expression profile of the REST and CoREST protein family in the human nervous system is not well understood and has been largely based on animal studies (Fuentes et al., 2012; Sáez et al., 2015; Wang et al., 2016). This gap in knowledge is important to address since genome wide binding profile studies of REST in the post-natal human and mouse hippocampus have shown that there are a number of unique binding sites in humans [22]. Similarly, Rockowitz and Zheng (23) showed that REST target sites were twice as many in human embryonic stem cells (ESCs) compared to mouse ESCs via chromatin immunoprecipitation coupled with sequencing (ChIP-seq) analysis (*n* = 8199 vs *n* = 4107). These findings suggest that the expression patterns of REST and CoREST proteins may vary significantly between species. Examining the expression profile of the CoREST family will provide insight into the cell types or developmental stages, at which they are most likely to elicit a function and the regulatory networks involved in orchestrating neurodevelopment. Thus, human models of neurogenesis, such as human pluripotent stem cells (hPSCs) should be employed to further interrogate the molecular changes involved in neurodevelopment.

This study aimed to use different 2D and 3D hPSC models to replicate human embryonic neurodevelopment and define a novel expression profile for *REST* and *RCOR* genes. The hPSCs were differentiated into glutamatergic and GABAergic forebrain neurons, using dual-SMAD inhibition, to allow for comparison of the expression levels in cortical neuronal subtypes. Transdifferentiation of hPSCs using NGN2 overexpression is routinely used to rapidly generate functionally mature, glutamatergic cortical neurons (as reviewed in Hulme, et al. (24)) and was utilised to explore *REST* and *RCOR* gene expression in differentiated neurons. SOX9 and NFIB overexpression was used to generate mature and functional astrocytes from hPSCs to define the levels of *REST* and *RCOR* genes in astrocytes. Cerebral organoids were differentiated from hPSCs as they are self-organised 3D structures that more closely resemble *in vivo* neurodevelopment. Understanding the expression pattern of *REST* and *RCOR* genes provides valuable insight into the cell types and stages during human neurodevelopment in which these transcription factors function and will then enable further research into identifying the target genes and molecular networks regulated by the CoREST family.

## Results

### *REST* and *RCOR* gene expression in 2D models of glutamatergic and GABAergic forebrain neurons derived from hESCs

To define an expression profile for *REST* and *RCOR* genes hESCs were differentiated into dorsal-derived forebrain glutamatergic neurons and ventral-derived forebrain GABAergic neurons using the dual-SMAD inhibition protocol (Figure 1A). Characterisation of neuronal gene markers in undifferentiated hESCs, glutamatergic and GABAergic hESC-derived forebrain neurons was completed by Nanostring nCounter, with findings presented as number of mRNA molecules normalised to a panel of 10 housekeeper genes (Figure S1A). Results were further supported by immunocytochemistry and RT-qPCR analyses (*n* = 3-4 independent differentiations; Figure S1B and C). The pluripotency marker *NANOG*, was highly expressed in hESCs (2703 mRNA molecules), with low levels in cultures differentiated to glutamatergic neurons (7 mRNA molecules) or GABAergic neurons (18 mRNA molecules) which indicates that cells have converted from the pluripotent state. *NKX2-1*, the gene that encodes a transcription factor critical in development of ventral forebrain neural progenitors, was shown to be only expressed in cultures differentiated to GABAergic neurons (1577 mRNA molecules). The cortical neural progenitor marker, *PAX6*, is highly expressed in cultures fated to glutamatergic neurons (1804 mRNA molecules). This was further supported by RT-qPCR analyses, showing a significant increase of *PAX6* expression in glutamatergic-fated neuronal cultures relative to hESCs (4958-fold change, *p* < 0.05; **Figure S1C**).

**Figure 1.**
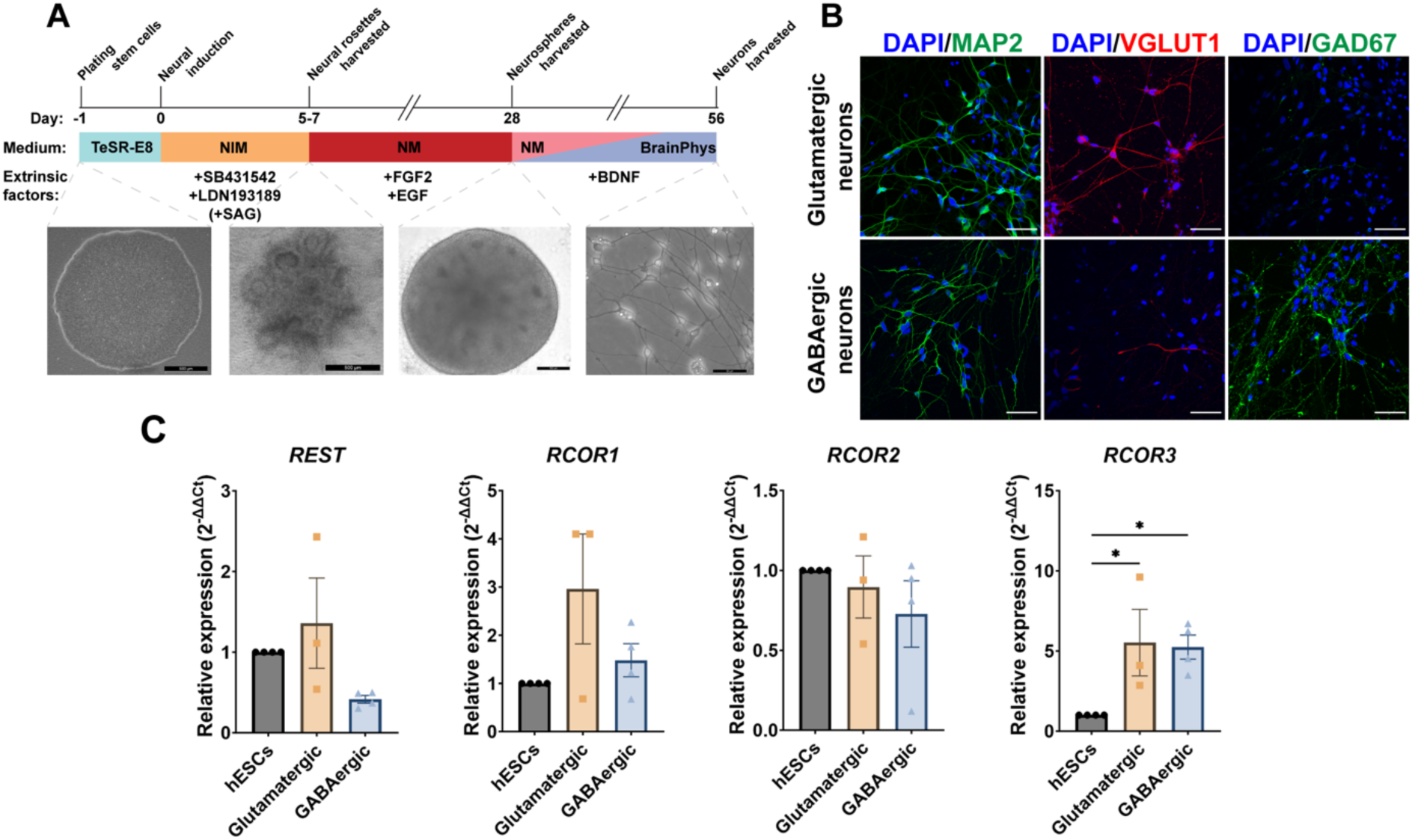
Expression profile of *REST* and *RCOR* genes in cortical and ventral forebrain neurons differentiated from human ESCs using dual-SMAD inhibition. **(A)** Schematic of differentiation protocol for glutamatergic and GABAergic neurons from hESCs. **(B)** Immunocytochemistry showing expression of MAP2 (green) in cultures fated to glutamatergic or GABAergic neurons, VGLUT1 (red) expressed predominantly in glutamatergic cultures and GAD67 (green) highly expressed in GABAergic cultures, nuclei are stained with DAPI (blue) (n = 3-4 independent differentiations). Scale bars = 50 µm. (C) *REST* and *RCOR* gene expression analysed as relative expression levels via RT-qPCR. Relative expression was calculated as the mean of three housekeeping genes and presented as mean ± SEM. Data was analysed using an ordinary One-way ANOVA with a Holm-Sidak test to correct for multiple comparisons. \**p* < 0.05.

High mRNA expression levels of general neuronal markers *GNG3*, *GRIA1/2*, *GRIN1*, *L1CAM*, *MAP2*, *MAPT*, *NDRG4*, *PSD95*, *SNAP91*, *SYN1* and *TUBB3* were identified in cultures fated to both neuronal cell types analysed by Nanostring (for protein names view Supplementary Table 1). The mature neuronal marker, microtubule associated protein-2 (MAP2) was expressed in both glutamatergic and GABAergic neuronal cultures, as confirmed by immunocytochemistry **(Figure 1A).** *MAP2* expression was also significantly increased in glutamatergic cultures (153-fold change, *p* < 0.05) and GABAergic cultures (138-fold change, *p* < 0.05) compared to hESCs by RT-qPCR **(Figure S1B).** The marker for cortical layer VI neurons, *TBR1* was expressed at high levels only in glutamatergic cultures and was significantly increased compared to both hESCs and GABAergic cultures (*p* < 0.05; **Figure S1BC)**. Vesicular glutamate transporter 1 (VGLUT1) a glutamate transporter found in glutamatergic neuronal populations, was expressed abundantly in glutamatergic cultures, with little expression in GABAergic cultures by immunocytochemistry **(Figure 1B)**. GABAergic, interneuron specific markers *DLX1/2*, *GAD2* and *SOX1* were solely expressed in GABAergic neuronal cultures detected by Nanostring. *GAD1*, the gene that encodes GAD67 a glutamic acid decarboxylase which functions in the synthesis of GABA, was shown to increase 240-fold in GABAergic cultures compared to hESCs analysed by RT-qPCR (*p* < 0.05; Figure S1B). GAD67 was also shown to be predominantly expressed only in GABAergic cultures via immunocytochemistry **(Figure 1A)**. There was little expression of cholinergic neuronal marker, *CHAT*, and dopaminergic neuronal marker, *TH*, suggestive of a lack of cholinergic and dopaminergic neuronal subtypes in either culture types. There were low levels of the glial progenitor marker *S100B* in neurons, suggesting a mixed population of neuronal and glial cells. There were decreased relative expression levels of the astrocyte marker, *GFAP*, in both neuronal culture types, compared to hESCs (*p* > 0.05). Collectively, the neuronal cultures were confirmed to express markers for glutamatergic and GABAergic forebrain neurons at the mRNA level via Nanostring and RT-qPCR and the presence of protein through immunocytochemistry, with low levels of astrocyte markers.

*REST* and *RCOR* gene expression levels were analysed in glutamatergic and GABAergic neuronal cultures using RT-qPCR to analyse relative fold changes **(Figure 1C)**. There was no significant difference in relative *REST* mRNA levels, however *REST* was decreased by 59 % in GABAergic cultures, compared to hESCs (*p* > 0.05). In addition, there was no significant difference observed in *RCOR1* mRNA expression levels, however glutamatergic cultures had a 3-fold increase and GABAergic cultures had a 1.5-fold increase in *RCOR1* mRNA. RT-qPCR analysis of *RCOR2* revealed both neuronal culture types remained similar to ESCs. *RCOR3* expression significantly increased 5.5-fold in glutamatergic cultures (*p* = 0.0445) and 5.2-fold in GABAergic cultures (*p* = 0.0421), when compared to hESCs, analysed by RT-qPCR. There were no significant differences in *REST* and *RCOR* gene expression between both neuronal culture types. Taken together, there was high variability in the expression pattern of *REST*, *RCOR1* and *RCOR2* in the glutamatergic and GABAergic neuronal cultures by dual-SMAD inhibition. This can be attributed to the high variability of differentiations and the heterogeneous neuronal cultures consisting of progenitors, glial progenitor cells and neurons. However, despite the variability in the fold change values, we obtained consistent results showing an increase in *RCOR3* expression upon neuronal differentiation of both hESC-derived cultures of glutamatergic and GABAergic neurons.

### *REST* and *RCOR* expression pattern in NGN2*-*induced neurons

Previous studies have described that induced expression of NGN2 rapidly promotes hPSC differentiation to functionally mature neurons making it a useful 2D model system for studying differentiated human neurons [as reviewed by 24]. We therefore utilised this approach to generate functional cortical neurons from three independent hPSC lines (H9 human ESCs (H9), iPSC1 and iPSC2 lines) to investigate expression of *REST* and *RCOR* genes as an alternative 2D model of human neurogenesis **(Figure 2A)**. Gene expression of neuronal markers in NGN2 induced neurons (iNs) were validated by RT-qPCR **(Figure S2)**. There was a significant increase for the mature neuronal marker *MAP2* in neurons derived from H9 (73.0- fold, *p* < 0.001), iPSC1 (40.3-fold, *p* < 0.05) and iPSC2 (31.9-fold, *p* < 0.05) cell lines. The layer VI cortical neuronal marker, *TBR1,* was also significantly increased in all three cell lines (H9 *p* < 0.001; iPSC1 *p* < 0.05; iPSC2 *p* < 0.05). *SLC17A7*, the gene that encodes the glutamatergic neuronal marker VGLUT1, was increased 3.8-fold in neurons derived from H9 stem cells, 30.4- fold in iPSC1 (*p* < 0.05) and 14.1-fold in iPSC2. *GAD1* (encodes for the GABAergic neuronal marker GAD67) was slightly increased in H9 and iPSC1 iNs (2.7- and 5.8-fold increase, respectively) and was decreased in iPSC2 iNs (44% reduction). In addition to neuronal marker gene upregulation, the cortical neural progenitor marker, *PAX6*, was increased across the three different hPSC lines (122.2-fold in H9 NGN2 iNs, 375.9-fold in iPSC1 and 16.3-fold in iPSC2), compared to undifferentiated stem cells. In NGN2 iNs from all 3 cell lines the astrocyte marker, *GFAP*, was not detectable after 40 cycles. Further validation of NGN2 iN differentiation was shown by immunocytochemistry, demonstrating protein expression of MAP2 and VGLUT1 **(Figure 2B)**. Finally, functional validation of NGN2 iNs was performed by electrophysiological analyses. The iNs derived from the H9, iPSC1 and iPSC2 cell lines recorded under current clamp displayed resting membrane potentials of - 55.6 ± 1.9 mV (*n* = 16), - 54.4 ± 2.5 mV (*n* = 12) and - 54.4 ± 1.6 mV (*n* = 16), respectively, consistent with the generation of functionally mature neurons. Figure 7C displays representative recordings of the three differentiated cell lines showing that the neurons generated are functionally active. Accordingly, negative current injections (−100 pA to −50 pA) elicited membrane hyperpolarizing responses, consistent with robust activity of hyperpolarization cyclic nucleotide gated potassium channels (HCN); whilst positive current injections (10 pA to 100 pA) evidenced robust multiple action potential firing **(Figure 2C)** typical of functional cortical neurons. Taken together, NGN2 iNs were functionally active and expressed mature cortical neuronal markers.

**Figure 2.**
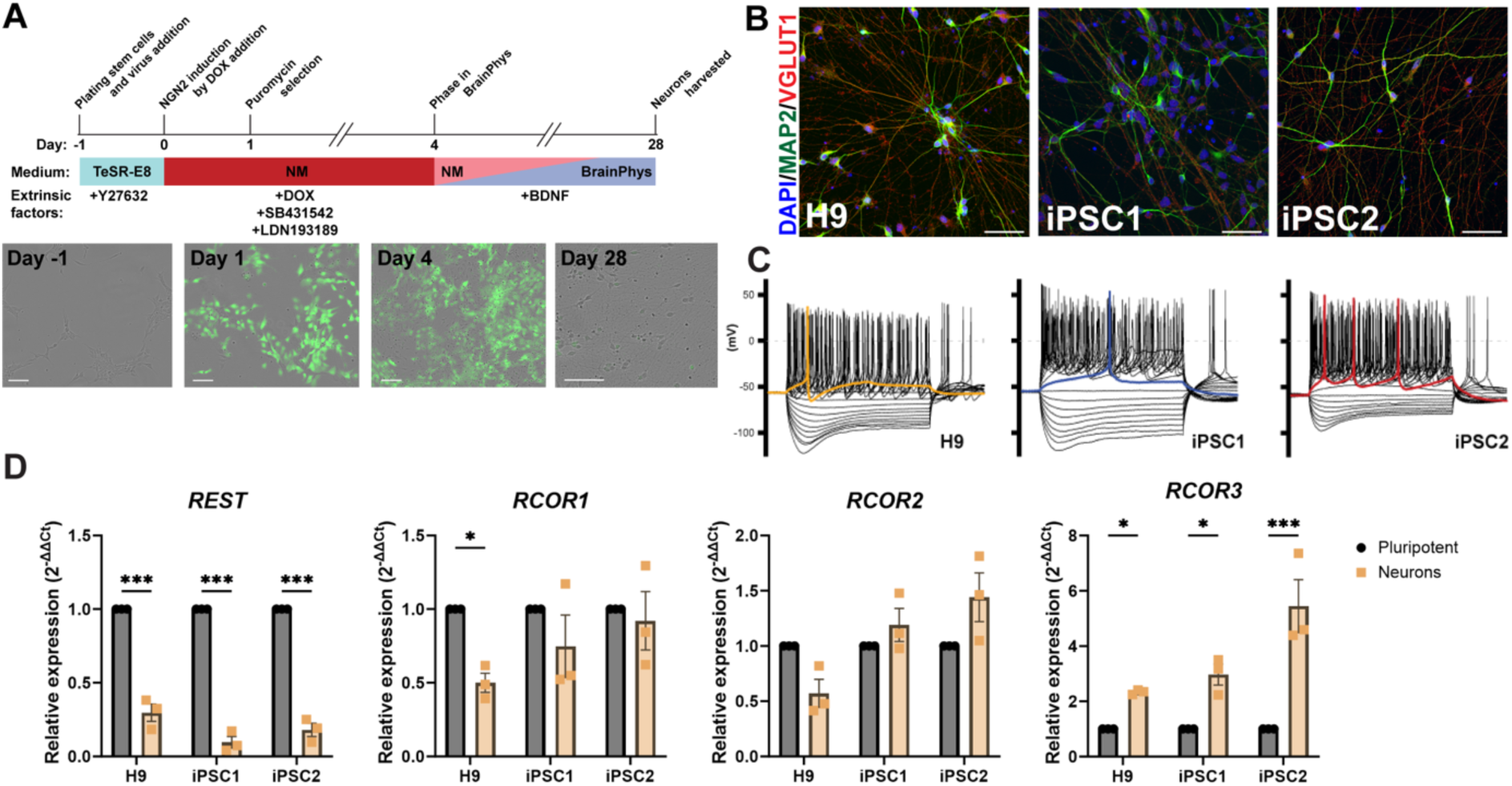
*REST* and *RCOR* gene expression in functional and mature NGN2 iNs. **(A)** hPSCs were differentiated into functionally mature iNs derived from H9 ESCs and two healthy control iPSC lines (iPSC1 and iPSC2) using NGN2 overexpression. **(B**) Immunocytochemistry confirmed expression of MAP2 and VGLUT1 at a protein level in neurons derived from all cell lines (n = 3 independent differentiations). Scale bars = 50 µm. **(C)** Representative current clamp recordings of membrane potential responses of the H9, iPSC1 and iPSC2 cell lines (n = 16, 12 and 16 cells, respectively, from 3 independent differentiations). The stimulation protocols consisted of 1 s long current injections from −100 pA to 100 pA in 10 pA steps. The coloured traces indicate firing activity at rheobase. **(D)** *REST* and *RCOR* gene relative expression levels were analysed via RT-qPCR. Data was calculated as the mean of three housekeeping genes and presented as mean ± SEM. Data was analysed using an ordinary One-way ANOVA with a Holm-Sidak test to correct for multiple comparisons. \**p* < 0.05, \*\*\**p* < 0.001. n = 3 biological replicates, with each data point representing the average of 3 technical replicates.

Following confirmation that NGN2 iNs expressed cortical neuronal markers and were functionally active, *REST* and *RCOR* gene expression was analysed via RT-qPCR **(Figure 2D)**. *REST* expression was significantly reduced in neurons derived from all cell lines when compared to hPSC (H9 71 %, iPSC1 93 % and iPSC2 82 %, *p* < 0.001). *RCOR1* was significantly decreased by 51 % only in H9 NGN2 iNs (*p* < 0.05). *RCOR2* was also decreased in H9-derived iNs by 43 % and remained unchanged in iNs derived from iPSC1 and iPSC2. Similar to neuronal cultures generated by dual-SMAD inhibition, *RCOR3* was shown to have an opposite expression profile to *REST* in NGN2 iNs. Expression was increased 2.3-fold in iNs derived from H9 (*p* < 0.05), 3-fold in iPSC1-derived (*p* <0.05) and 5.4-fold in iPSC2-derived (*p* < 0.001) iNs. The iNs derived from iPSC2 had the lowest level of *PAX6* expression and also the highest expression of *RCOR3* compared to iNs derived from the other cell lines. Consistent with all models *REST* was shown to decrease during neuronal differentiation, whereas *RCOR3* had the opposite expression profile and increased with neuronal differentiation.

### CoREST3 exhibits nuclear subcellular localisation in neurons

The subcellular localisation of CoREST3 in neuronal differentiation was characterised using immunocytochemistry and the % CoREST3 positive cells were calculated in hESCs and neuronal subtypes **(Figure 3)**. In undifferentiated hESCs, CoREST3 displayed nuclear expression and colocalised with the pluripotent marker, OCT4. CoREST3 also showed nuclear localisation in cultures of glutamatergic and GABAergic neurons derived from H9 hESC using the dual SMAD approach **(Figure 3A**). There was no significant difference in the % CoREST3 positive cells between cell types and neuronal differentiation methods **(Figure 3B)**. These findings suggest that CoREST3 is ubiquitously expressed in cortical neuronal subtypes and is localised in the nucleus.

**Figure 3.**
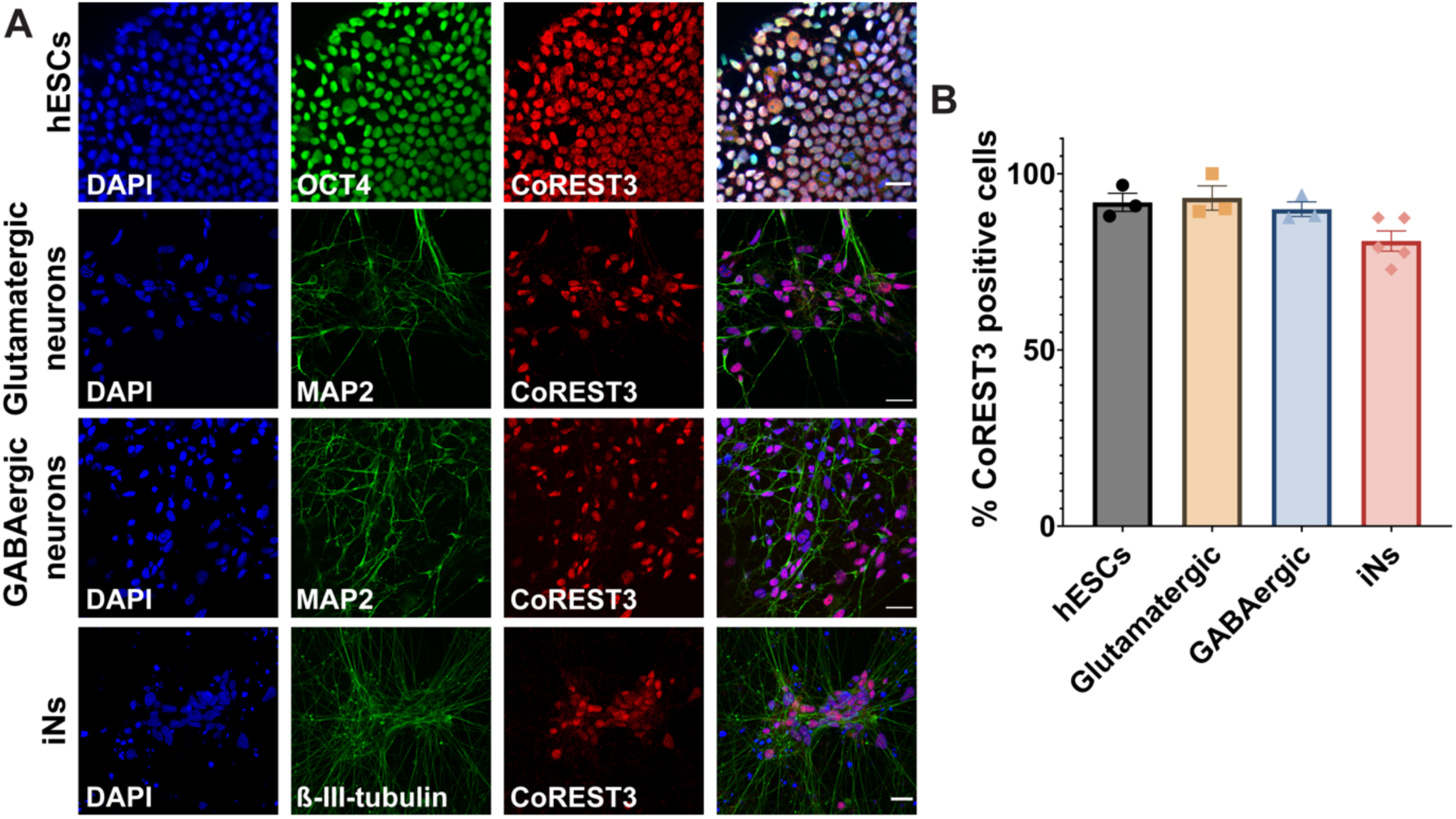
CoREST3 exhibits nuclear subcellular localisation and is ubiquitously expressed in different neuronal subtypes. **(A)** Immunocytochemistry was used to determine the subcellular localisation of CoREST3 in H9 hESCs, H9-derived neuronal cultures of glutamatergic, GABAergic and iNs. CoREST3 was shown to be expressed in the nucleus of all cell types. Scale bar = 50 µm. **(B)** The % CoREST3 positive cells were calculated over the total number of nuclei and across neuronal differentiation and subtypes (n = 3-5 fields of view, n = 3-5 independent differentiations).

### *REST* and *RCOR* gene expression in hPSC-derived astrocytes

To define the expression profile of *REST* and *RCOR* genes in human astrocytes, we firstly adapted a transdifferentiation protocol, published by Canals et al. (33), which generated mature functional astrocytes in 21 days using inducible expression of the mouse genes, *Nfib* (encodes Nuclear factor 1B) and *Sox9* (encodes SRY Box transcription factor 9). Our study further streamlined this protocol by using a single inducible vector to ectopically express human *NFIB* and *SOX9* **(Figure 4A).** The induced astrocytes (iAs) generated by *SOX9* and *NFIB* overexpression were characterised by immunocytochemistry and RT-qPCR to confirm the expression of astrocyte markers. The pan-astrocyte marker, GFAP, was expressed in the cytoplasm of astrocytes at both day 7 and day 14 **(Figure 4B).** There were 70.7%, 72.3% and 67.1% GFAP+ astrocytes derived from H9, iPSC1 and iPSC2 cell lines, respectively at day 14. GFAP was significantly increased in day 14 astrocytes derived from the iPSC2 cell line (*p* < 0.05) (Figure 4C). The iAs also expressed astrocyte and glial markers at the protein level, including NFIB, SOX9, S100ß, EAAT2 and ALDOC **(Figure S3).** To further validate astrocytes generated by SOX9 and NFIB overexpression, one-step RT-qPCR was used to confirm expression of general astrocyte markers at day 0 (hPSCs), and day 7 and 14 of maturation **(Figure S4A).** *S100ß* was significantly increased after 7- and 14-days maturation in H9 and iPSC1 iAs (*p* < 0.01). There were no significant differences in expression of *S100ß* between day 7 and 14 of maturation in all three lines. The iAs matured for 14 days had a 16.2-fold, 122.7- fold, 218.5-fold increase in *GFAP* expression compared to undifferentiated hPSCs in H9 (*p* > 0.05), iPSC1 (*p* < 0.001) and iPSC2 (*p* < 0.001) cell lines, respectively. *SLC1A3* (encoding EAAT1) was significantly increased at both day 7 and 14 for all three cell lines (*p* < 0.05), whereas *SLC1A2* (encoding EAAT2) was only significantly increased in day 14 iAs in all three cell lines (*p* < 0.05). There was no increase in expression of neuronal markers *TUBB3* and *MAP2* in day 14 iAs compared to hPSCs in all three cell lines, analysed by RT-qPCR (Figure S4B). The iAs cultures had significantly lower expression of both neuronal genes, compared to neurons (*p* < 0.05), and there was no difference in expression levels between hPSCs and iAs. Together, these findings suggest the population of iAs are a pure population of astrocytes, with no evidence of neurons present in the culture.

**Figure 4.**
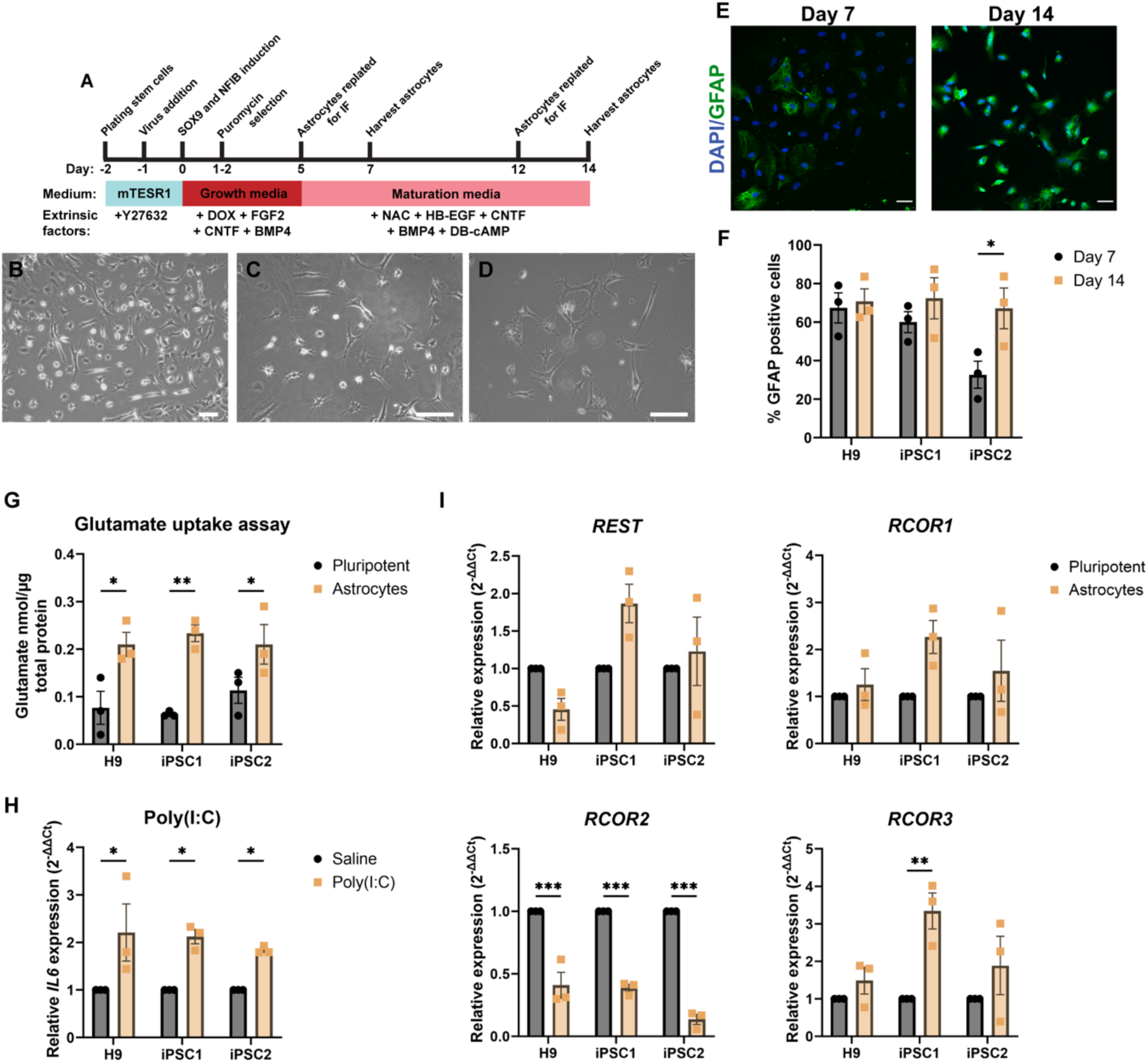
Expression profile of *REST* and *RCOR* genes in human astrocytes. **(A)** Flow diagram for induced astrocyte generation. Representative images of iAs after 7 days **(B),** 14 days **(C)** and 21 days **(D)** of maturation. Scale bar = 100 µm. **(E)** Representative images of iAs stained for astrocyte marker, GFAP, at 7 and 14 days of maturation from iPSC2 cell line (Scale bar = 50 µm), **(F)** and the % positive cells for each target were calculated. Each data point represents the average of 3-4 fields of view per independent differentiation (n = 3 independent differentiations). **(G)** A glutamate uptake assay was completed on day 14 iAs derived from H9, iPSC1 and iPSC2 cell lines and determined by a serial dilution of glutamate standard (samples run in duplicate from 3 independent differentiations). **(H)** Day 14 iAs were incubated with Poly(I:C) and *IL6* (encodes Interleukin-6) expression was analysed using one-step RT-qPCR to assess the ability to respond to inflammatory stimulation. **(I)** The expression levels of *REST* and *RCOR* genes was assessed in day 14 iAs derived from three hPSC cell lines using RT-qPCR. n = 3 independent differentiations, n = 3 technical replicates. n = 3 independent differentiations, n = 3 technical replicates. Data was analysed using a Two-way ANOVA with a Holm-Sidak test for multiple comparisons, \**p* < 0.05, \*\**p* < 0.01, \*\*\**p* < 0.001

Finally, assessments of glutamate uptake and inflammatory activation were performed to functionally validate iAs. There was a significant increase in glutamate uptake in day 14 iAs, compared to hPSC in all three cell lines (∼ 2-fold, *p* < 0.05)**(Figure 4G)**. Furthermore, following incubation with Poly(I:C), iAs had a significant increase in *IL6* (encodes Interleukin-6) expression across all three cell lines **(Figure 4h)**. Taken together, these findings demonstrate that iAs have the ability to uptake glutamate and respond to inflammatory stimulation.

Gene expression analysis of *REST* and *RCOR* genes were performed on iAs after 14 days of maturation using RT-qPCR **(Figure 4I).** There were no significant differences in *REST* and *RCOR1* expression between hPSCs and iAs for all three lines (*p* < 0.05). *RCOR2* was significantly decreased in iAs by 59.0%, 61.4% and 86.4% in H9, iPSC1 and iPSC2 cell lines, respectively (*p* < 0.001), whereas *RCOR3* was significantly increased 3.3-fold in iAs derived from iPSC1 (*p* < 0.01), with no significant differences in iAs generated from H9 or iPSC2 cell lines. The subcellular localisation of CoREST3 was assessed using immunocytochemistry, with representative images of iAs derived from iPSC2, shown in **Figure S5A.** CoREST3 was shown to be expressed ubiquitously in GFAP-positive iAs and appeared in both the nucleus and cytoplasm. To further validate the subcellular localisation of CoREST3, different secondary antibodies were used to support the observations **(Figure S5B).** Together, these findings highlight the distinct expression profile of *REST* and *RCOR* genes and the subcellular localisation of CoREST3 in human astrocytes.

### *REST* and *RCOR* genes display differential expression patterns in human astrocytes and neurons

To compare the expression levels of *REST* and *RCOR* genes in hPSC-derived astrocytes and neurons, gene expression analysis was performed on day 14 iAs and day 21 iNs using RT-qPCR **(Figure 5).** *REST* was significantly decreased in iNs derived from iPSC1 and iPSC2 compared to iAs (94.9%, *p* < 0.001 and 85.4%, *p* < 0.05, respectively). There were no significant changes in *REST* mRNA levels in iNs and iAs derived from the H9 stem cell line. There was a trend for lower *RCOR1* expression levels in iNs derived from all three cell lines, compared to iAs, with a significant reduction in iNs from iPSC1 (67.0%, *p* < 0.05). There was a 3.1-fold increase in *RCOR2* expression in iNs derived from iPSC1 (*p* < 0.01) and a 10.6-fold increase in iNs derived from iPSC2 (*p* < 0.001) compared to iAs. There were no significant differences in *RCOR3* expression between iNs and iAs derived from H9 and iPSC1 lines, however there was a 2.9- fold increase in iNs compared to iAs derived from iPSC2 (*p* < 0.01). These results suggest there are cell type dependent expression profiles for *REST* and *RCOR* genes in human neurodevelopment.

**Figure 5.**
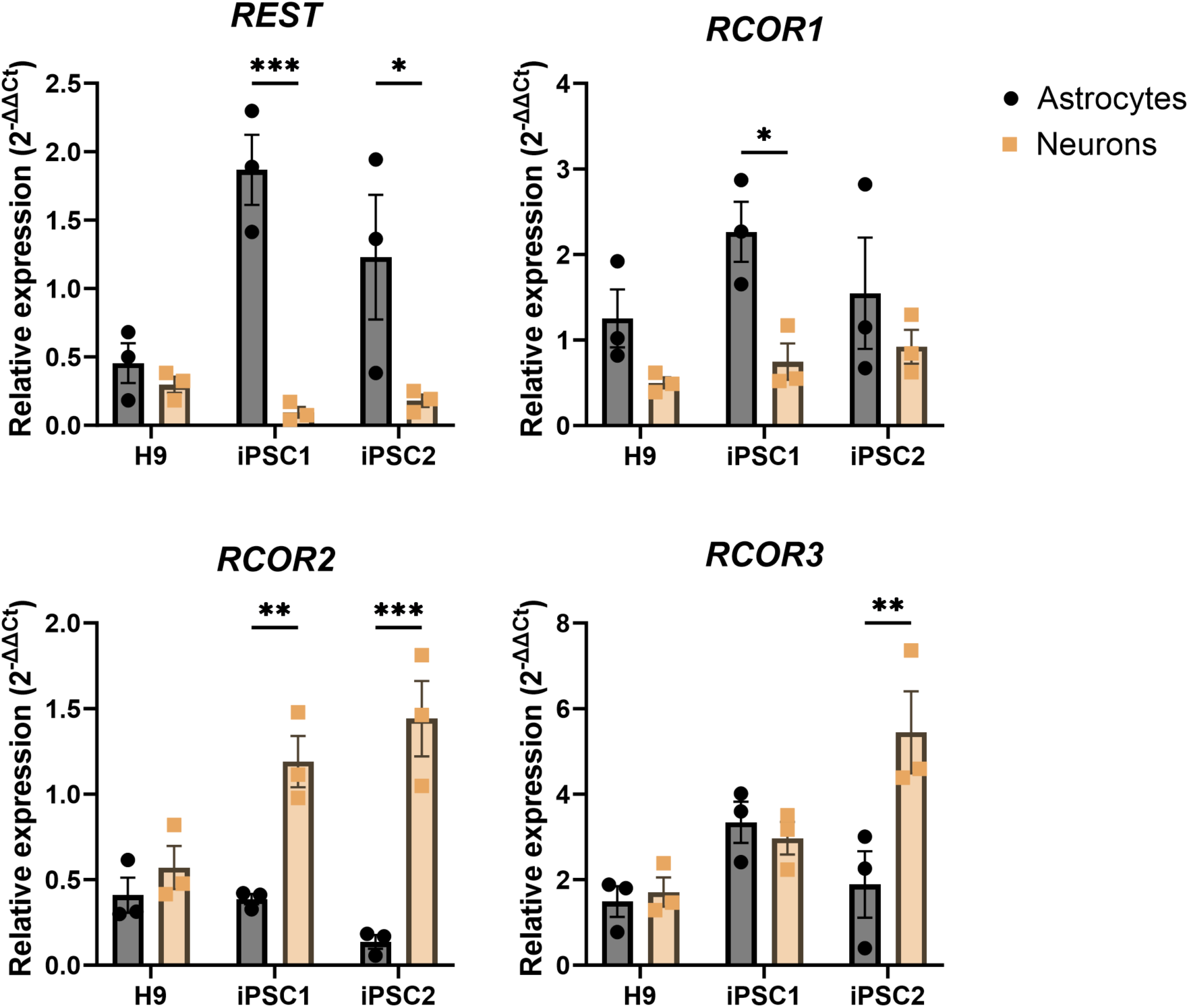
*REST* and *RCOR* genes have different expression levels in human astrocytes and neurons. RT-qPCR was performed on day 14 iAs generated by SOX9-NFIB overexpression and day 21 iNs generated by NGN2 overexpression to compare differences in mRNA levels of *REST* and *RCOR* genes. n = 3 independent differentiations, n = 3 technical replicates. Data is presented as fold changes compared to hPSC and analysed using Two-way ANOVA with a Holm-Sidak test for multiple comparisons.

### *REST* and *RCOR* expression pattern in 9 month-matured cerebral organoids

To further explore the expression profile of *REST* and *RCOR* genes in human neurodevelopment, 9 month-matured cerebral organoids differentiated from two healthy control iPSC lines (iPSC1 and iPSC2) were utilised as a 3D tissue model **(Figure 6A)**[50]. Organoids derived from iPSC1 and iPSC2 displayed expansion and maturation monitored via organoid area increase over the first three months **(Figure 6B),** and expressed MAP2 and GFAP confirmed at the protein level via immunocytochemistry **(Figure 6C).** Cerebral organoids were characterised by neuronal and glial gene expression patterns via RT-qPCR **(Figure S6).** The mature neuronal marker, *MAP2*, was shown to increase significantly in iPSC1 (38.7-fold increase, *p* < 0.001) and iPSC2 (13.9-fold increase, *p* > 0.05) organoids compared to undifferentiated iPSCs. Organoids were shown to have a proportion of cells destined to differentiate into glutamatergic neurons evidenced by a significant increase of *TBR1* expression in iPSC1 and in iPSC2 (*p* < 0.05), compared to undifferentiated iPSCs. *SLC17A7*, the gene that encodes VGLUT1, was shown to increase 10.4-fold in both iPSC1 and 10.3-fold in iPSC2 organoids (*p* < 0.05). Cerebral organoids also presented expression of *GAD1,* a marker representative of GABAergic interneurons in iPSC1 (46.2-fold change, *p* < 0.01) and iPSC2 (8.6- fold change, *p* > 0.05), compared to undifferentiated iPSCs. The astrocyte marker, *GFAP*, was increased 78.5-fold in iPSC1 (*p* > 0.05) and 656.9-fold increase in iPSC2 (*p* > 0.05) organoids. Taken together, these results suggest the cerebral organoids differentiated from two iPSC lines are a mixed population of glutamatergic and GABAergic neuronal subtypes and glial cells, specifically astrocytes.

**Figure 6.**
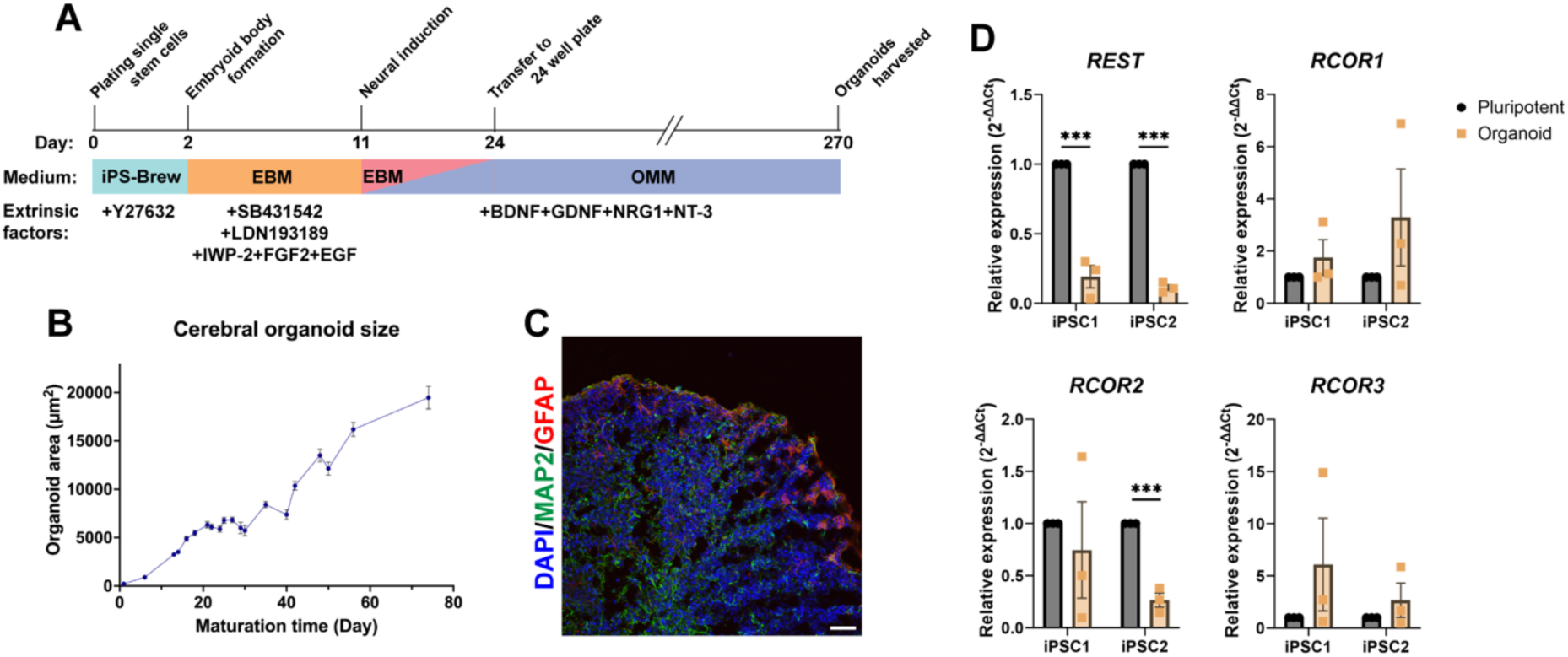
Expression pattern of *REST* and *RCOR* genes in human cerebral organoids derived from two healthy control iPSC lines (iPSC1 and iPSC2). **(A)** Cerebral organoids were matured for 9 months prior to harvesting. **(B)** Organoids were shown to be a mixed population of neurons (MAP2+) and astrocytes (GFAP+) via immunocytochemistry. **(C)** Organoid area (µm^2^) was measured over time with a steady increase in organoid size observed over maturation time. (D) *REST* and *RCOR* genes expression levels were analysed using RT-qPCR. Relative expression is calculated from the mean of three housekeeping genes and presented as mean ± SEM. Data was analysed with a Two-way ANOVA with statistical significance determined using the Holm-Sidak multiple comparison method. \**p* < 0.05, \*\**p* < 0.01, *** *p* < 0.001.

Following characterisation, *REST* and *RCOR* gene expression was analysed in cerebral organoids **(Figure 6D).** Reflective of the expression of mature neuronal markers, *REST* mRNA levels were significantly reduced in both iPSC1 (81 %, *p* < 0.001) and iPSC2 (89 %, *p* < 0.001) organoids, compared to undifferentiated iPSCs. *RCOR1* levels increased 1.7-fold and 3.3-fold in organoids derived from iPSC1 and iPSC2 (*p* > 0.05). Similar to *REST, RCOR2* expression decreased by 25 % (*p* > 0.05) and 74 % (*p* < 0.001) in iPSC1 and iPSC2 organoids, respectively. Although not significant, there was a trend for increased *RCOR3* expression by 6.1-fold and 2.7-fold in iPSC1 and iPSC2, respectively (*p* > 0.05). In summary, organoids were shown to be a mixed population of neuronal subtypes and glial cells, with *REST* and *RCOR* gene expression differences in these co-cultures potentially influenced by cell type ratio differences. The expression trends from this data are, however, consistent with the data recorded in 2D monocultures of hPSC-derived forebrain neurons, highlighting the distinct expression pattern of *REST* and *RCOR* genes, suggestive of potential unique functions.

## Discussion

This study is the first to provide the expression profile for *REST* and *RCOR* genes in human neural development using 2D and 3D hPSC models. *REST* and *RCOR2* levels decreased in neurons and organoids, *RCOR3* was upregulated in neurons and *RCOR2* transcripts were downregulated in astrocytes. The expression profile provides valuable insight into the cell types in which these transcription factors may be eliciting a function and will aid in deepening our understanding of the molecular mechanisms regulated by the CoREST family.

Dynamic expression patterns of *REST* and *RCOR* genes are indicative of distinct functions in different neural cell types, and provide insight into when these transcription factors are likely to elicit a function. Current literature is suggestive of distinct expression patterns of each CoREST protein during neurodevelopment and in the mature brain depending on the cell type and developmental stage (as reviewed in Maksour, et al. (15)). Our knowledge of the expression changes of the REST and CoREST family is largely based on rodent studies, with differences observed in REST target genes between species [23,40]. This study aimed to identify the expression pattern for *REST* and *RCOR* genes in human neurogenesis using 2D and 3D hPSC models. *REST* levels were shown to decrease in more mature models of neuronal differentiation supporting the notion that REST expression decreases to allow expression of neuronal genes, as seen in multiple models [41,42,43]. *REST* expression was only decreased in more functional neuronal cultures, highlighting the importance of using multiple models and validating these to support the findings. *RCOR1* expression was shown to be significantly reduced in the mature NGN2-induced functional neurons, suggesting CoREST1 maintains a function at the immature neuronal stage that is lost with neuronal maturation. This is in-line with data from the embryonic developing mouse cortex, where CoREST1 expression increased until post-natal day 15 before decreasing in the adult cortex [17]. *RCOR2* expression was shown to be significantly decreased in cerebral organoids and induced astrocytes, compared to undifferentiated stem cells, a novel finding suggesting CoREST2 may play a role in repressing astrocyte or glial genes whereby a decrease in expression may allow for glial differentiation. This finding is supported in spontaneous senescence-accelerated P8 mouse astrocytes, showing *Rcor2* expression was significantly decreased compared to controls and possibly responsible for regulating neuroinflammation through the interplay with *IL6* [20]. Future studies that perform gene knockdown of *RCOR2* in neural progenitor cells are needed to observe whether this drives towards an astrocyte lineage, in addition to *RCOR2* gene knockdown in astrocytes and comparing changes in inflammatory activation. Interestingly, *RCOR3* exhibited a distinct expression profile to the other paralogues, with expression increasing in neurons and astrocytes derived from hPSCs. This result is suggestive that CoREST3 may play a unique role in mature neurons and astrocytes, compared to the CoREST family. The higher expression of *RCOR3* in neurons is supported by freely available datasets from the Brain RNA-seq, which shows *REST, RCOR1* and *RCOR2* levels to be lower than *RCOR3* in neurons [44] (**Figure S7**). The nuclear subcellular localisation of CoREST3 in neurons is consistent with the known role of CoREST3 as a regulator of gene expression, whereas the cytoplasmic expression in astrocytes may suggest that either CoREST3 has additional, yet unknown roles in the cytoplasm, or that subcellular localisation may regulate CoREST3 functions in astrocytes. Gene editing of *RCOR3* in mouse models or human derived neural cells will provide insight into the function of CoREST3 in the brain.

Cerebral organoids provide a platform to better replicate physiological development, compared to 2D cell cultures, as they are self-assembled structures composed of different neuronal subtypes, glial cells and progenitors that not only mimic the interactions of the developing brain at a cellular level but also in terms of tissue architecture and functional neuronal networks [29,45,46,47,48,49,50]. The cerebral organoid data in this study highlighted a downregulation of *REST,* in line with mature neurons, a reduction in *RCOR2,* concomitant with astrocytes, and a trend for *RCOR3* upregulation, in line with neurons and astrocytes. The expression patterns were shown to have high variability across replicates and cell lines, suggesting the composition of a mixture of neural cells of differing maturity. The heterogeneous nature of the organoids may explain why subtle changes are not observed for each gene, as was the case in monocultures. Additionally, transcriptomic analysis of brain organoids has highlighted that individual organoids replicate endogenous brain development by distinct and indistinguishable cellular diversity and network formations accompanied by oscillatory activity patterns that are similar to those observed in preterm human electroencephalography [51,52,53]. The organoid-to-organoid variability observed explains the spread of data seen in the *RCOR* gene expression analysis, as organoids are a diverse cell population of progenitors, neuronal subtypes, and glial cells all of different maturation stages. Therefore, this study demonstrates the importance of using monocultures for expression profiling, and organoids to act as an accessible substitute to developing brain tissue samples.

### Conclusions

In summary, this study provided the first expression profile for *REST* and *RCOR* genes in human neural development using 2D and 3D hPSC models. Similar patterns were observed across multiple models of hPSC neuronal differentiation, whereby *REST*, *RCOR1* and *RCOR2* levels decreased with neuronal maturation. This was contrasted with *RCOR3,* which showed upregulated expression with hPSC neuronal differentiation. Increased expression of *RCOR3* was also observed in hPSC astrocyte differentiation, together with an increased expression of *RCOR2*. Overall, the dynamic expression patterns of *REST* and *RCOR* genes observed during hPSC neuronal and glial differentiation highlights the potential distinct functions that REST and CoREST proteins may play in regulating development of these cell types in humans.

## Materials and Methods

### Cell culture and cell lines

Use of hPSC lines for this thesis project was approved by UOW Human Ethics committee (#2017-375, 2017-382, 2020-450, 2020-451, 13-299). The H9 human embryonic stem cells (ESCs) line (WA09 obtained from WiCell; referred to as ESC) and two healthy control induced pluripotent stem cell (iPSC) lines, iPSC1 [25] and iPSC2 [26] were utilised in the current study. hPSCs were cultured as previously reported in Abu-Bonsrah, et al. (27), Denham and Dottori (28) and Mattei, et al. (29). Briefly, the hPSCs were maintained in TesR-E8 (StemCell Technologies, #05990) or mTESR1 (StemCell Technologies, #85850) on vitronectin XF (StemCell Technologies, #7180) coated tissue culture ware, kept in normoxic conditions at 37 °C with 5% CO_2_. Cells were passaged once every 5-7 days using 0.5 mM EDTA (Life Technologies, #AM9260G) in phosphate-buffered saline (PBS)^-/-^ (Life Technologies, #14190250).

### Differentiation of hESCs into cortical and ventral forebrain neurons using dual-SMAD inhibition

Forebrain neurons were generated using a modified protocol as published in Denham and Dottori (28) and Nasr, et al. (30)(Figure 1A). Briefly, hESC colonies are passaged and small clusters are plated onto a 6 well plate at a density of ∼ 1-2 x 10^4^ cells/well in TeSR-E8 media (StemCell Technologies, #5990). Following 24 hrs, media was removed and neural induction media (NIM; Supplementary Table 2) supplemented with small molecules 10 µM SB431542 (StemCell Technologies, #72146) and 0.1 µM LDN193189 (StemCell Technologies, #72234) and additionally 0.4 µM SAG (Sigma-Aldrich, #566660) for ventral forebrain differentiation) was added, with full media changes every 2^nd^ day. Following neural induction for 5-7 days, neural rosettes were harvested by 0.5 mM EDTA (in PBS^-/-^) dissociation and gently tapping the side of the culture plate or pipetting to dislodge 3D neural rosette-like structures. Cells were centrifuged at 200 x *g* for 2 min and resuspended in neural media (NM; Supplementary Table 2) supplemented with 0.02 µg/mL FGF2 (StemCell Technologies, #78003) and 0.02 µg/mL EGF (StemCell Technologies, #78006.1). To each well, 100 µL was plated into an ultra-low attachment U-bottom 96 well plate using a multi-channel pipette, and plates were centrifuged at 200 x *g* for 2 min to bring aggregates together. Neurospheres began to form between 18 and 24 hrs after plating. A top up media change of 50 µL of NM with supplements was completed 3 days after plating out. Half media changes with NM and growth factors were continued every 3^rd^ day. After 2-3 weeks, neurospheres were collected into a 15 mL falcon tube and old media removed. Accutase (ThermoFisher Scientific, #00-4555-56) was added to the neurospheres for 10 min at 37 °C, fresh media was added to inactivate the enzyme and cells pelleted at 200 x *g* for 2 min. Single cells were resuspended and counted. Approximately 7 x 10^5^ cells/well were plated onto each well of 10 µg/mL Poly-D-lysine (PDL, Sigma-Aldrich, P6407-5MG; coated for 30 min at RT and washed twice with PBS) and 10 µg/mL laminin (LAM, ThermoFisher Scientific, #23017015; coated overnight at 4 °C) coated 12 well plates in NM supplemented with 10 ug/mL BDNF (StemCell Technologies, #78005). Neurons were observed after 24-72 h. Half media changes were completed every second day. To improve maturation, BrainPhys media (Supplementary Table 2) was subsequently added at increasing concentrations (25-100% in NM) for each media change. After 5-7 days of neuronal maturation (or when non-neuronal cells were apparent), cytosine arabinoside (AraC; Sigma Aldrich, #C1768) was added for 48 hrs to purify the neuronal culture from dividing cells. LAM was supplemented once a week in the media at 1 ug/mL to promote attachment if cells appeared to be lifting off from the plate. Neurons were harvested after 4 weeks of maturation.

### Lentiviral production

Viral particles containing an open reading frame of *Neurogenin-2* (*NGN2*) were produced to differentiate hPSCs into mature neurons or an open reading frame of *SRY-box9* (*SOX9*) and *Nuclear Factor I B* (*NFIB*) to generate astrocytes. Briefly, HEK293T cells were transfected with the DNA of lentiviral packaging plasmids vSVG (Addgene, USA, #8454), RSV (Addgene, #12253), pMDL (Addgene, #12251), and either the tetracycline transactivator (TTA) vector, M2rtTA (Addgene, #20342), the *NGN2* overexpression vector, TetO-*NGN2*-eGFP-Puro plasmid (Addgene, #79823), or one of the *RCOR3*-shRNA-mCherry-BSD plasmids or the TetO-*SOX9*-*NFIB*-mPlum-Puro plasmid using Polyethyleneimine (Sigma-Aldrich, USA, #408727). DNA is added in a ratio of 4:2:1:1, transfer vector:pMDL:RSV:vSVG. The cell culture media containing viral particles was collected every 24 h over 3 days. The viral supernatant was concentrated by ultracentrifugation at 66,000 x *g* for 2 hours at 4 °C. Viral pellet was resuspended in PBS and stored at −80°C until needed.

### Generation of NGN2 induced neurons (iNs)

This study used a modified version of the protocol published by Zhang, et al. (31) and Nehme, et al. (32) to generate mature neurons via NGN2 overexpression (Figure 2A). Briefly, hPSCs are resuspended as single cells using accutase for 2 min at RT following a passage. Single cells are plated at 10,000 cells/cm^2^ onto 10 µg/mL PDL and LAM coated culture plate in TeSR-E8 media supplemented with 10 µM Y27632. Cells are allowed to attach for 6-8 h, after which 0.5 - 1 µL of viral particles of both NGN2 overexpression and the TTA per well. Virus was removed 16-20 h following transduction with fresh NM (Supplementary Table 1) supplemented with 1 µg/mL doxycycline (DOX; Sigma-Aldrich, # D9891), 10 µM SB431542 and 0.1 µM LDN193189 to promote a cortical fate. After 24 h of DOX induction, 0.5 µg/mL puromycin was added daily for 3 days for selection of successfully transduced cells, in addition to DOX, SB431542 and LDN193189. Following selection fresh media supplemented with 10 µg/mL BDNF was added. Following selection, BrainPhys media (Supplementary Table 1) was subsequently added at increasing concentrations (25-100% in NM) for each media change to improve maturation. Neurons were matured for 21-28 days after viral transduction prior to being harvested.

### Generation of induced astrocytes (iAs) by SOX9 and NFIB overexpression

Astrocytes were generated using a modified protocol based on Canals, et al. (33) and summarised in **Figure 3A**. Briefly, 6 well plates were coated with Matrigel (Bio-strategy, #BDAA354277) diluted 1:200 in base medium for 30 min at 37 °C. Following standard passaging and plating for maintenance, hPSCs were centrifuged at 300 x *g* for 3 min and resuspended in 500 µL accutase (ThermoFisher Scientific, #00-4555-56) for 3 min at RT. Accutase was inactivated with 1 mL fresh mTESR1 media and pelleted at 300 x *g* for 3 min and resuspended in fresh mTESR1 supplemented with 10 µM Y27632 media. Single cells were plated out at 100,000 cells/well in fresh mTESR1 media supplemented with 10 µM Y27632. Cells were allowed to attach for 6 - 8 h, after which 5 µL of viral particles of both the *SOX9/NFIB* overexpression and the TTA vector per well was added directly to each well. Virus was removed the following day with a full media change of growth media (GM; Supplementary Table 2) supplemented with 1 µg/mL DOX. After 24 hrs of DOX induction, 2 µg/mL puromycin was added for 48 h to select successfully transduced cells, in fresh GM addition to DOX. Following selection, a full media change with GM supplemented with DOX was performed. 5 days after SOX9 and NFIB activation, GM media was transitioned to maturation medium (MM; Supplementary Table 2) for the life of the culture, with half media changes performed every second day. To compare the differences in astrocyte differentiation with different mediums, media was changed to commercial astrocyte growth supplement media (SM) (Supplementary Table 2) on day 10 for the duration of the culture, with half media changes performed every other day. On day 5 and 12, astrocytes were disassociated with accutase and plated out into black wall optical 96 well plates (Corning, #3603) at a cell density of 10,000 cells/well for immunocytochemistry and functional assays were performed on day 7 and 14 astrocytes. Astrocytes were harvested at day 7 (**Figure 3B**), day 14 (**Figure 3C**) and day 21 (**Figure 3D**) for One-step RT-qPCR.

### Generation of cerebral organoids from two healthy control iPSC lines

Cerebral organoids were generated using a modified protocol based on Salick, et al. (34). Briefly, iPSC colonies were passaged using 0.5 mM EDTA (in PBS^-/-^) as near single cell suspension, plated onto Matrigel (Corning, #354277) coated 60 mm cell culture dishes (5000 cells per dish) and maintained in TeSR-E8 media for five days in hypoxic conditions (3% O2, 5% CO2, 37°C). Five day old iPSC colonies were collected via PBS-EDTA (ThermoFisher Scientific, #14190-144, #15575-038), dissociated into a single cell suspension, and seeded at 1000 cells per well into U-bottom 96 well plates (Corning, #7007) in iPS-Brew (Miltenyi Biotec, #130-107-086) containing 10 µM Y27632 (ROCK inhibitor; Calbiochem, #688000), followed by brief centrifugation (30 s at 100 x *g*) to focus the cell suspensions. Partial media changes were performed every 48 h. Embryoid body media (EBM) was used for days 2 to 11 (Supplementary Table 2) and supplemented with 10 µM SB431542 (Stemcell Technologies, #72234), 0.1 µM LDN193189 (MedChemExpress, #HY-12071) and 1 µM IWP-2 (MedChemExpress, #HY-13912) until day 8, and 20 ng/µL FGF2 (Stemcell Technologies, #130-104-923) and 20 ng/µL EGF (Miltenyi Biotec, #130097750) until day 11. Neural induction was triggered by phasing in organoid maturation media (OMM) (Supplementary Table 2) with partial media changes every 48 h until day 24. Spheroids where then transferred to 24 well plates to accommodate the larger media volume needs, maintained in normoxic conditions (20% O_2_, 5% CO_2_, 37 °C), and matured through the addition of 5 ng/mL BDNF (Miltenyi Biotec, #130-096-286) and 10 ng/mL GDNF (Miltenyi Biotec, #130-098-449) from day 26, and 10 ng/mL NRG1 (Peprotech, #AF-100-03) and 20 ng/mL NT-3 (Miltenyi Biotec, #130-093-973) from day 34, after which partial media changes were performed every 3 to 4 days until the end of the 9 months (∼270 days).

### Gene quantification

#### RT-qPCR

RT-qPCR analysis was completed as described in Ng, et al. (35). Briefly, total RNA was harvested from cell cultures using the PureLink RNA Mini Kit (Life Technologies, #12183025) as per manufacturer’s instructions. RNA quality was assessed using the NanoDrop 2000 Spectrophotometer (ThermoFisher Scientific, USA) with an A_260/280_ ratio ranging from 1.9 to 2.1 deemed acceptable. Up to 1000 ng of RNA was used to synthesise cDNA using the iScript™ gDNA Clear cDNA Synthesis Kit (Bio-Rad, USA, #1725035) following the manufacturer’s guidelines. PowerUp™ SYBR™ Green Master Mix (ThermoFisher Scientific, #A25778) was used for the RT-qPCR step. Primers were used at 400 nM concentration and are displayed in Supplementary Table 3. Primer annealing temperatures were optimised by a serial dilution standard curve and considered acceptable within a range of 85-110% efficiency. Each reaction was run in triplicate and contained 20 ng of cDNA template in a final reaction volume of 20 µL. Cycling parameters were: 50 °C for 2 min for UDG activation, 95 °C for 2 min, then 40 cycles of 95 °C for 1 s and the annealing/extending temperature for each primer displayed in Supplementary Table 3 for 30 seconds, followed by the melt curve stage of 95 °C for 15 seconds, 60 °C for 1 min and 95 °C for 15 seconds. RT-qPCR reaction and data collection was performed using the QuantStudio 5 Real-Time PCR System (Applied Biosystems, USA) and raw data exported to Microsoft Excel for analysis. ΔCt values were obtained by normalisation to the average of three house keeper genes, *β2M, GAPDH* and *PPIA*. ΔΔCt values were obtained by normalisation to ΔCt values of control samples. Using the 2^-ΔΔCt^ method relative gene expression values (fold change) was determined. Results were presented as the mean ± SEM.

#### One-step RT-qPCR

One-step RT-qPCR was performed as described in [36]. RNA was extracted by adding TBST (5 mM Tris, 75 mM NaCl and 0.05% Triton X-100) directly to cells for 5 min. RT-qPCR reactions were carried out with 5 µL template volumes in a final reaction volume of 10 µL volumes in MicroAmp 96-Well Fast Reaction Plates (ThermoFisher, #4366932). Each target or multiplex combination was prepared with TaqPath 1-Step RT-qPCR Master Mix (ThermoFisher, A15300) as per the manufacturer’s guidelines. This study utilised 0.5 × master mix and predesigned assay for gene expression analysis was used without loss of amplification. Probes for *GFAP* (ThermoFisher, Hs00909233_m1), *S100B* (ThermoFisher, Hs00389217_m1), *SLC1A3* (encodes EAAT1; ThermoFisher, Hs00188193_m1), *SLC1A2* (encodes EAAT2; ThermoFisher, Hs01102423_m1), *IL6* (encodes Interleukin 6; ThermoFisher, Hs00174131_m1) and the housekeeping gene *TBP* (ThermoFisher, Hs00427620_m1). RT-qPCR reactions were performed on the Quantstudio 5 (ThermoFisher) with the following thermal cycling conditions; reverse transcriptase for 50 °C for 15 min, enzyme activation 2 min, amplification 95 °C for 1 s/60 °C for 15 s. Data was analysed with Quantstudio Design & Analysis Software v1.5.1 and exported to Microsoft Excel.

### Immunocytochemistry

Cell monolayers were fixed in 4% (w/v) paraformaldehyde at RT for 20 min and then washed twice in PBS. Samples were then permeabilised with 0.3% (v/v) Triton X-100 (Sigma-Aldrich, #T9284) in PBS for 10 min at RT. Blocking buffer [10% (v/v) normal donkey serum (Sigma-Aldrich, #D9663) in PBS] was used to block the samples for 1 hr at RT. Samples were subsequently incubated with primary antibodies (Supplementary Table 4) prepared in blocking buffer at 4°C overnight. Samples were then washed 3x with 0.1% (v/v) Triton X-100 in PBS and incubated with donkey-anti-mouse IgG (H+L) Alexa Fluor 488 nm (1:500, Abcam, UK, ab150109), donkey-anti-rabbit IgG (H+L) Alexa Fluor 555 nm (1:500, Abcam, ab150062) and donkey-anti-goat IgG (H+L) Alexa Fluor 647 nm (1:500, Abcam, ab150135) labelled secondary antibody conjugates prepared in blocking buffer for 1 hr at RT. Nuclei were stained using DAPI (1:500; Sigma-Aldrich, #D9542) in PBS at RT for 15 min. Prolong Gold antifade reagent (Invitrogen, USA, #P36934) was utilised to preserve fluorescence signal and mount samples onto glass slides. Samples were imaged with a Leica TSC SP8 II confocal microscope (Leica Microsystems, Australia). Fiji was used to collect and analyse images [37].

### Nanostring

A custom CodeSet was designed to be used with the PlexSet Nanostring technology (Nanostring) to analyse the genes in Supplementary Table 1. The mRNA from ESCs, cortical neurons and ventral neurons was extracted as described in Section 2.3.1. The mRNA concentration and the processing of the Nanostring run was performed as previously described in Muñoz, et al. (38).

### Electrophysiology

Plastic coverslips that were cut into rectangular slides smaller than 10 mm in width and coated with PDL and LAM as described above prior to plating NGN2 iNs onto the slides for functional characterisation. Whole-cell patch clamping was conducted on neurons that were matured between 3-4 weeks and as described by Hulme, et al. (39). Briefly, whole-cell patch clamp recordings were made at room temperature (20-22 °C) with a MultiClamp 700B Amplifier, digitalized with a Digidata 1440, and controlled with pClamp11 software (Molecular Devices; LLC, San Jose, CA, USA).

### Statistical analyses

Experiments were performed with at least 3-4 independent experimental differentiations (*n* = 3-4, biological replicates) and 3 technical replicates of each sample. Statistical analysis was conducted using GraphPad Prism software, version 8.0.2 (GraphPad Software, La Jolla, USA). Data was determined to be normally distributed using a Shapiro-Wilk normality test, if normally distributed, data was analysed by a one-way ANOVA with Tukey’s post hoc or multiple t-test with Holm-Sidak method, unless stated otherwise. An α of 0.05 (*p*-value < 0.05) was considered statistically significant.

## Acknowledgments

The authors would like to thank Dr Tracey Berg for input and advice into vector design and Dr Diane Ly for kindly donating polyethyleneimine. We thank Distinguished Professor David J. Adams, University of Wollongong, Illawarra Health & Medical Research Institute, for providing support towards the electrophysiology studies.

## Author Contributions

Conceptualization, S.M., L.O., and M.D.; methodology, S.M., N.N., A.J.H., Sara M., M.E., S.S.M., R.B., B.R. R.F.U.; formal analysis, S.M.; investigation, S.M., N.N., A.J.H., Sara M., M.E., S.S.M., R.B., B.R. R.F.U.; data curation, S.M.; writing—original draft preparation, S.M.; writing— review and editing, S.M., L.O., and M.D.; visualization, S.M.; M.E.; R.F.U.; supervision, L.O., and M.D.; project administration, S.M., L.O., and M.D. All authors have read and agreed to the published version of the manuscript.

## Ethics statement

All subjects gave their informed consent for inclusion in the study. The study was approved by the UOW Human Ethics committee (#2017-375, 2017-382, 2020-450, 2020-451, 13-299).

## Data Availability Statement

The raw data supporting the conclusions of this article will be made available by the corresponding author, without undue reservation.

---

**Figure S1. Characterisation of glutamatergic and GABAergic forebrain neuronal differentiations via dual-SMAD inhibition. (A)** Heatmap of the number of mRNA molecules for neuronal genes by a custom panel using the Nanostring nCounter (*n* = 1). (B) Neuronal gene expression in glutamatergic and GABAergic neuronal cultures were analysed by RT-qPCR analysis (*n* = 3-4 independent differentiations, *n* = 3 technical replicates). Relative expression is calculated to the mean of three housekeeping genes and presented as mean ± SEM. Data was analysed using a One-way ANOVA with Holm-Sidak test for multiple comparisons, if data was not normally distributed a Kruskal-Wallis test corrected for multiple comparisons using a Dunn’s test. \**p* < 0.05, \*\**p* < 0.01, *** *p* < 0.001. Gene details are listed in **Table S4**.

**Figure S2. Molecular characterisation of H9 human ESCs and two healthy control iPSC lines (iPSC1 and iPSC2) differentiated into induced neurons (iNs) via NGN2 overexpression.** Molecular analysis of NGN2 iNs was completed using RT-qPCR. Data is presented as the mean ± SEM from three independent differentiations (n = 3 biological replicates) with each data point representing the average of 3 technical replicates. Data was analysed using an ordinary One-way ANOVA with a Holm-Sidak test to correct for multiple comparisons. \**p* < 0.05, \*\*\**p* < 0.001.

**Figure S3. Immunocytochemistry images and analysis of astrocyte markers in induced astrocytes. (A)** Representative images of iAs stained for astrocyte markers at 7 and 14 days of maturation from iPSC2 cell line (Scale bar = 50 µm), **(B)** and the % positive cells for each target were calculated. Each data point represents the average of 3-4 fields of view per independent differentiation (n = 3 independent differentiations). Data was analysed using a Two-way ANOVA with a Holm-Sidak test for multiple comparisons, \**p* < 0.05, \*\**p* < 0.01, \*\*\**p* < 0.001.

**Figure S4. iAs express general astrocyte markers and do not express neuronal genes at an mRNA level. (A)** One-step qPCR of general astrocyte markers in hPSCs (Day 0), and after 7- and 14-days maturation revealed significant increases in *S100ß, GFAP, SLC1A3* (encodes EAAT1) and *SLC1A2* (encodes EAAT2) in iAs derived from the H9, iPSC1 and iPSC2 cell lines. **(B)** RT-qPCR of general neuronal markers, *TUBB3* (encodes ß-III-tubulin) and *MAP2*, showed no differences in expression levels between hPSCs and astrocytes, with NGN2 iNs having significantly higher expression of both markers in all three hPSC lines. n = 3-4 independent differentiations, n =3 technical replicates. Data was analysed with a Two-way ANOVA with Holm-Sidak test for multiple comparisons.

**Figure S5. Representative immunocytochemistry images of CoREST3 subcellular localisation in astrocytes. (A)** Representative immunocytochemistry images of iAs derived from iPSC2 are shown, with DAPI (blue), GFAP (green) and CoREST3 (red). Magnified images are highlighted by the white rectangle. Scale bar = 50 µm. **(B)** CoREST3 was stained with an anti-rabbit 488 nm (#A11008) and anti-rabbit 555 nm (#A21207) secondary antibodies to confirm that the same subcellular localisation was observed. Scale bar = 50 µm.

**Figure S6. Molecular characterisation of human cerebral organoids derived from two healthy control iPSC lines (iPSC1 and iPSC2).** Cerebral organoids were matured for 9 months prior to harvesting RNA and analysing gene expression profiles for neuronal and glial markers via RT-qPCR **(A-E)**. Relative expression is calculated from the mean of three housekeeping genes and presented as mean ± SEM. Data was analysed with a Two-way ANOVA with statistical significance determined using the Holm-Sidak method. \**p* < 0.05, \*\**p* < 0.01, *** *p* < 0.001.

**Figure S7. Cell type specific RNA-seq data of the expression profile of *REST* and *RCOR* genes in the human brain.** Data is sourced from the Brain RNA-seq database which is a collection of RNA-seq of purified cell types from human brains [44].

