## Supplementary materials for "*REST* and *RCOR* genes display distinct expression profiles in neurons and astrocytes using 2D and 3D human pluripotent stem cell models"

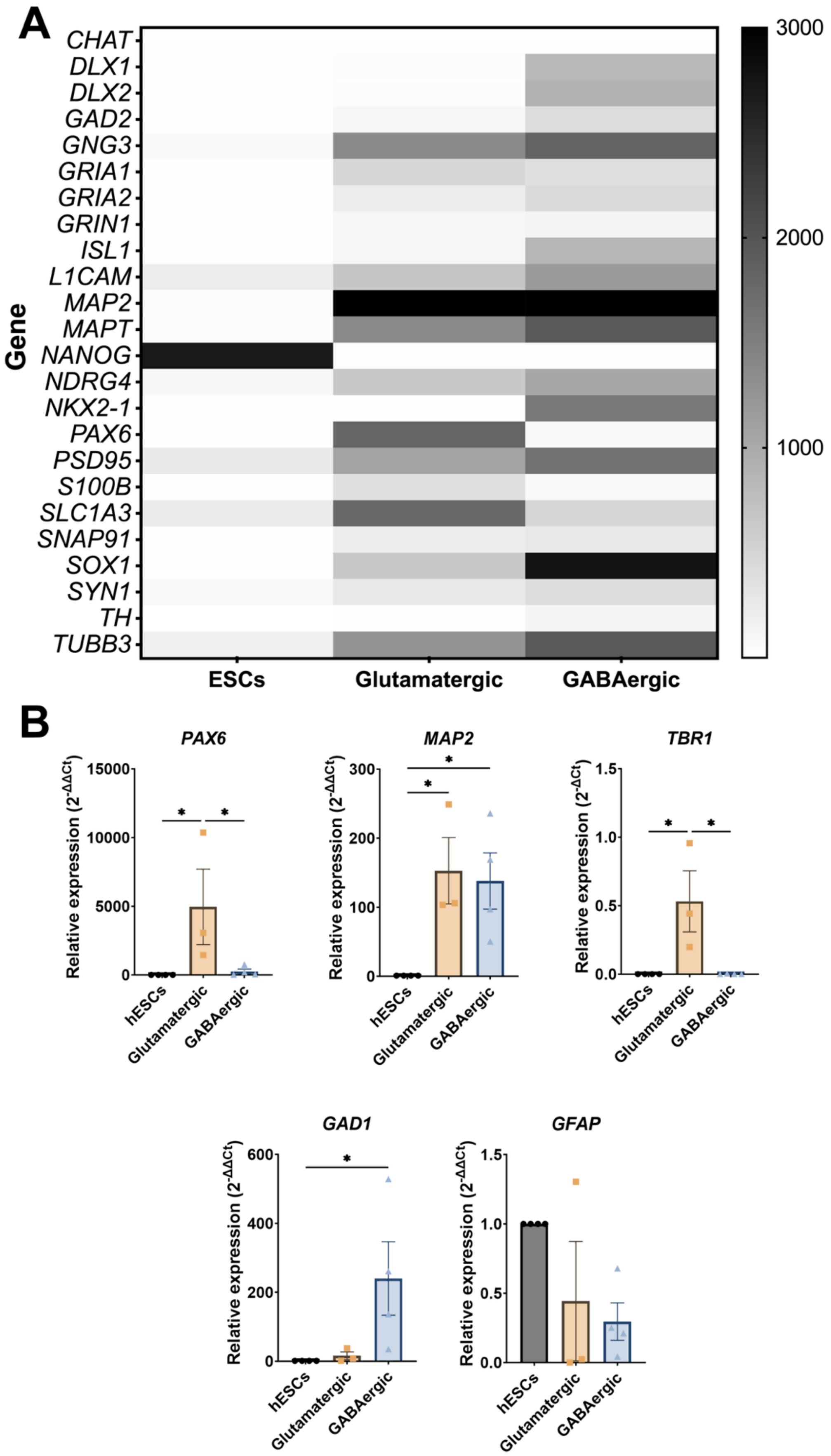

**Figure S1. Characterisation of glutamatergic and GABAergic forebrain neuronal differentiations via dual-SMAD inhibition. (A)** Heatmap of the number of mRNA molecules for neuronal genes by a custom panel using the Nanostring nCounter (*n* = 1). (B) Neuronal gene expression in glutamatergic and GABAergic neuronal cultures were analysed by RT-qPCR analysis (*n* = 3-4 independent differentiations, *n* = 3 technical replicates). Relative expression is calculated to the mean of three housekeeping genes and presented as mean ± SEM. Data was analysed using a One-way ANOVA with Holm-Sidak for multiple comparisons, if data was not normally distributed a Kruskal-Wallis test corrected for multiple comparisons using a Dunn's test. *$p < 0.05$, **$p < 0.01$, *** $p < 0.001$. Gene details are listed in **Table S4**.

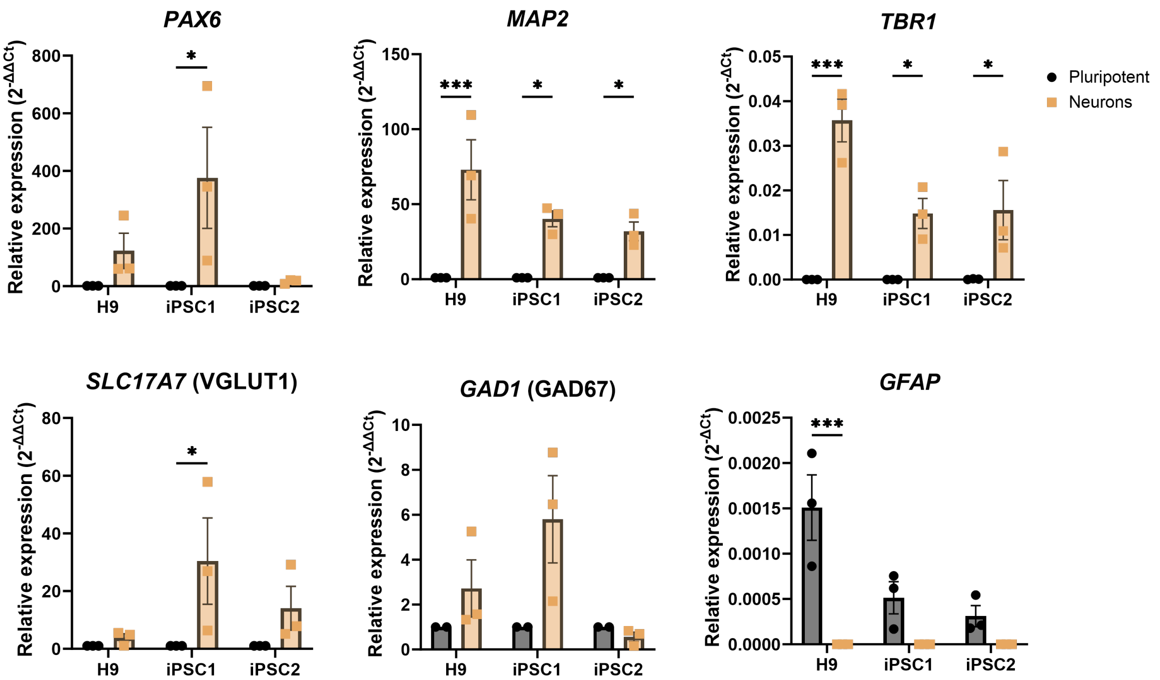


**Figure S2. Molecular characterisation of H9 human ESCs and two healthy control iPSC lines (iPSC1 and iPSC2) differentiated into induced neurons (iNs) via NGN2 overexpression.** Molecular analysis of NGN2 iNs was completed using RT-qPCR. Data is presented as the mean ± SEM from three independent differentiations (n = 3 biological replicates) with each data point representing the average of 3 technical replicates. Data was analysed using an ordinary One-way ANOVA with a Holm-Sidak to correct for multiple comparisons. *$p < 0.05$, ***$p < 0.001$.

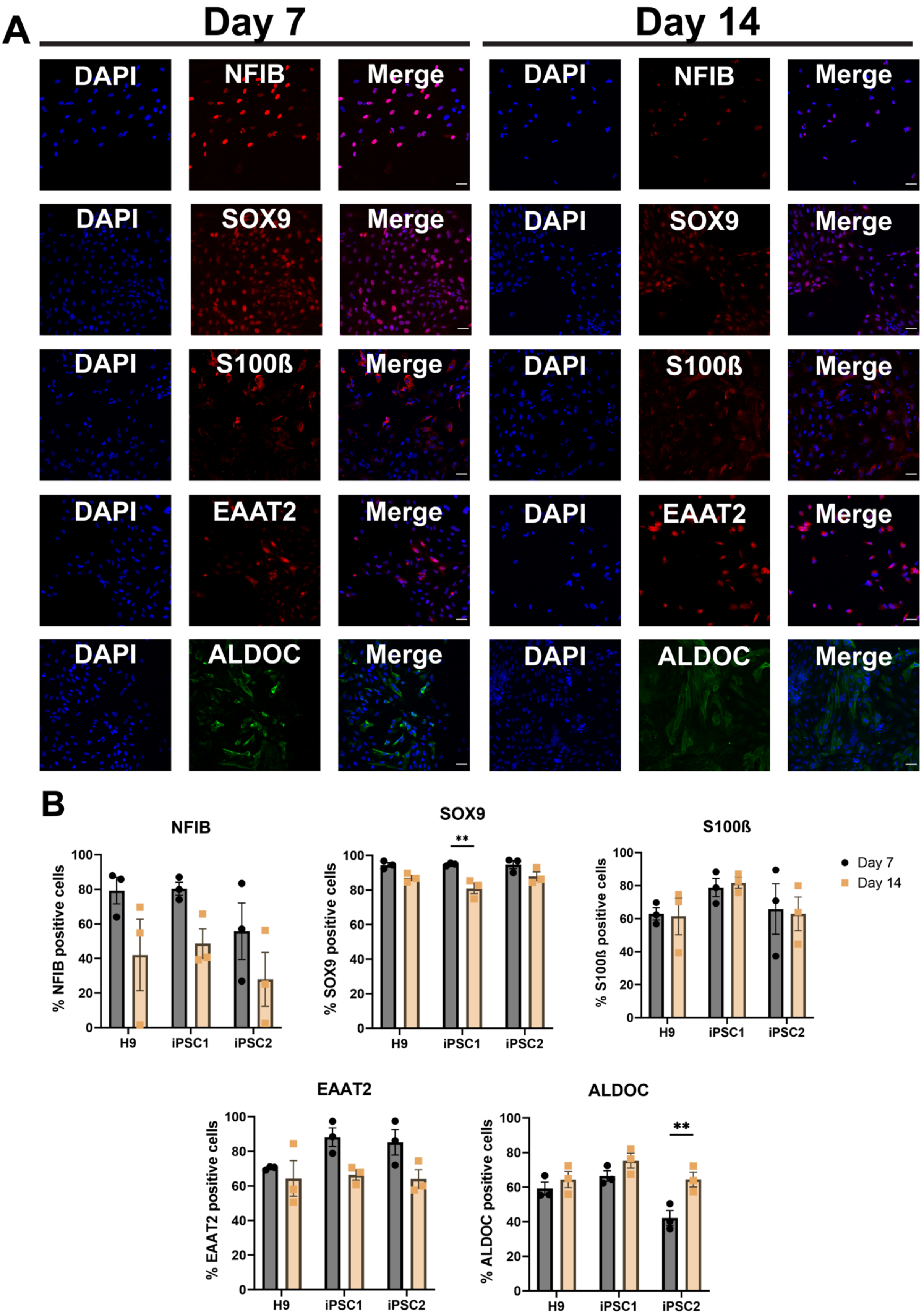


**Figure S3. Immunocytochemistry images and analysis of astrocyte markers in induced astrocytes. (A)** Representative images of iAs stained for astrocyte markers at 7 and 14 days of maturation from iPSC2 cell line (Scale bar = 50 µm), **(B)** and the % positive cells for each target were calculated. Each data point represents the average of 3-4 fields of view per independent differentiation (n = 3 independent differentiations). Data was analysed using a Two-way ANOVA with a Holm-Sidak for multiple comparisons, *$p < 0.05$, **$p < 0.01$, ***$p < 0.001$.

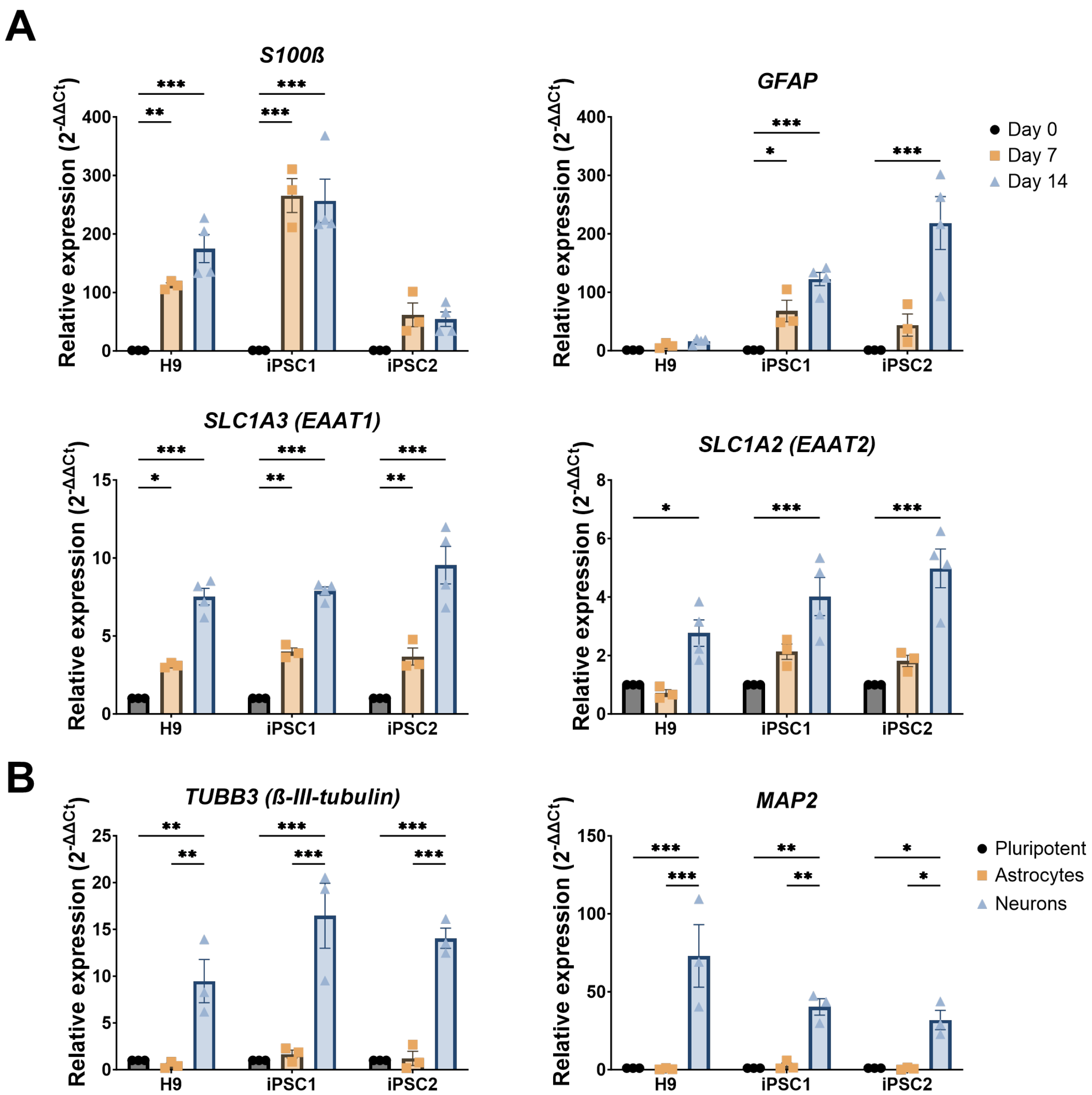


**Figure S4. iAs express general astrocyte markers and do not express neuronal genes at an mRNA level. (A)** One-step qPCR of general astrocyte markers in hPSCs (Day 0), and after 7- and 14-days maturation revealed significant increases in *S100ß, GFAP, SLC1A3* (encodes EAAT1) and *SLC1A2* (encodes EAAT2) in iAs derived from the H9, iPSC1 and iPSC2 cell lines. **(B)** RT-qPCR of general neuronal markers, *TUBB3* (encodes ß-III-tubulin) and *MAP2*, showed no differences in expression levels between hPSCs and astrocytes, with NGN2 iNs having significantly higher expression of both markers in all three hPSC lines. n = 3-4 independent differentiations, n =3 technical replicates. Data was analysed with a Two-way ANOVA with Holm-Sidak for multiple comparisons.

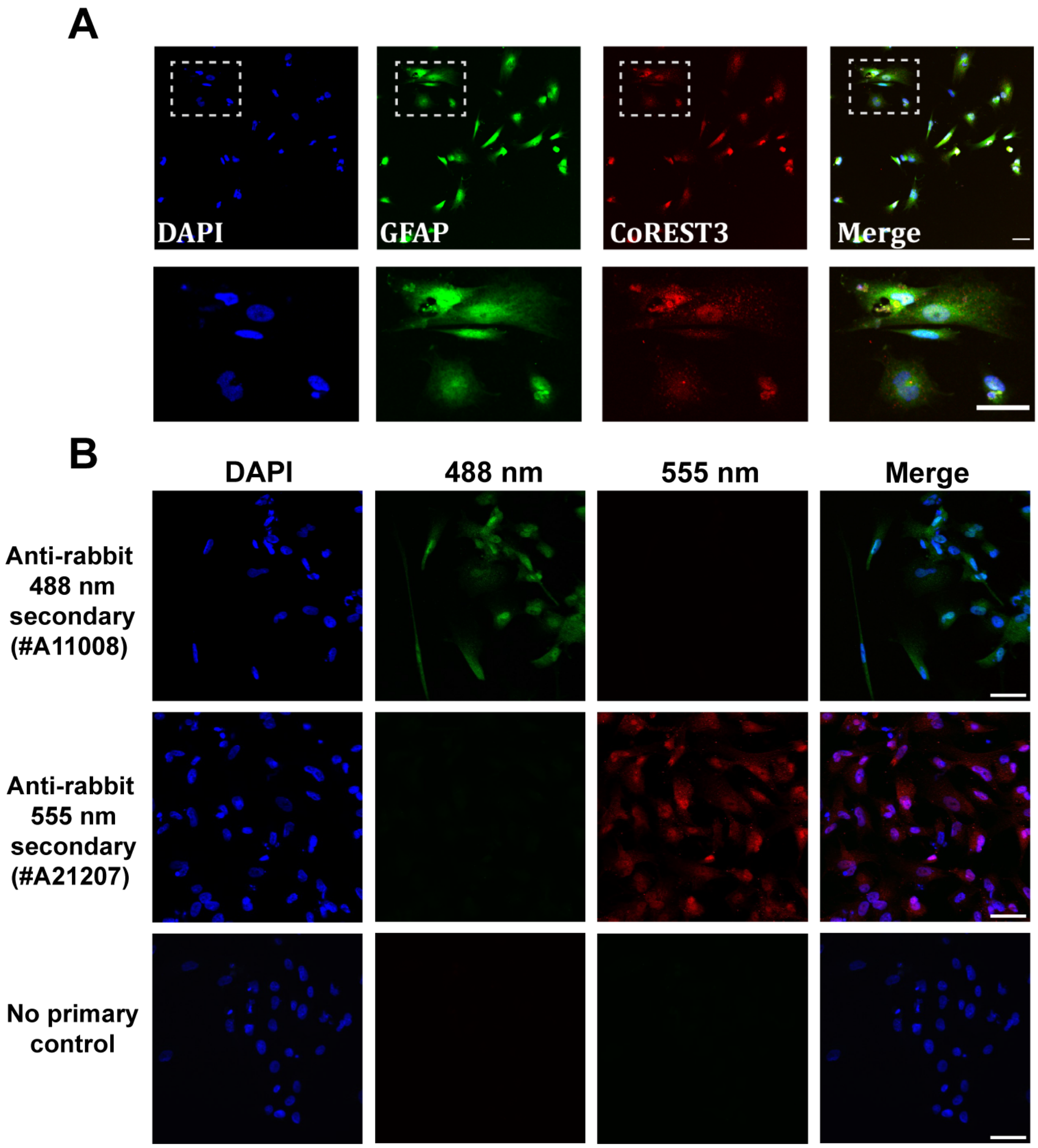


**Figure S5. Representative immunocytochemistry images of CoREST3 subcellular localisation in astrocytes.** **(A)** Representative immunocytochemistry images of iAs derived from iPSC2 are shown, with DAPI (blue), GFAP (green) and CoREST3 (red). Magnified images are highlighted by the white rectangle. Scale bar = 50 µm. **(B)** CoREST3 was stained with an anti-rabbit 488 nm (#A11008) and anti-rabbit 555 nm (#A21207) secondary antibodies to confirm that the same subcellular localisation was observed. Scale bar = 50 µm.

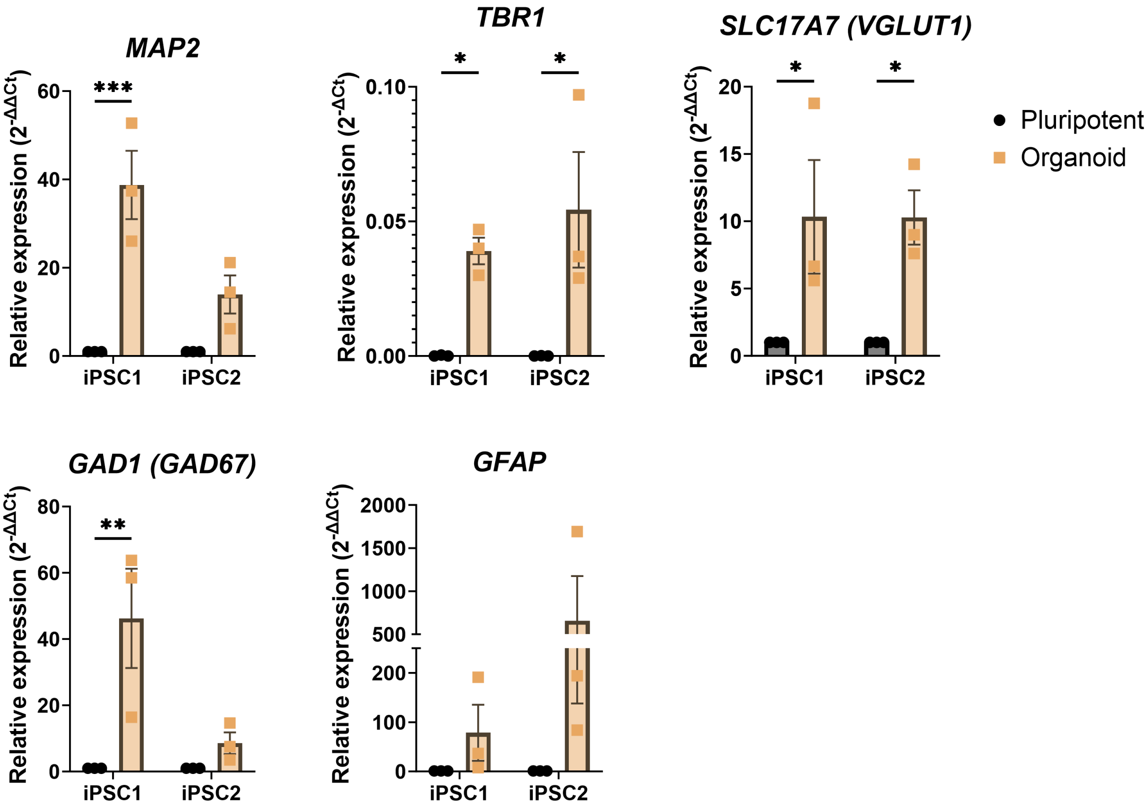


**Figure S6. Molecular characterisation of human cerebral organoids derived from two healthy control iPSC lines (iPSC1 and iPSC2).** Cerebral organoids were matured for 9 months prior to harvesting RNA and analysing gene expression profiles for neuronal and glial markers via RT-qPCR **(A-E)**. Relative expression is calculated from the mean of three housekeeping genes and presented as mean ± SEM. Data was analysed with a Two-way ANOVA with statistical significance determined using the Holm-Sidak method. *$p < 0.05$, **$p < 0.01$, *** $p < 0.001$.

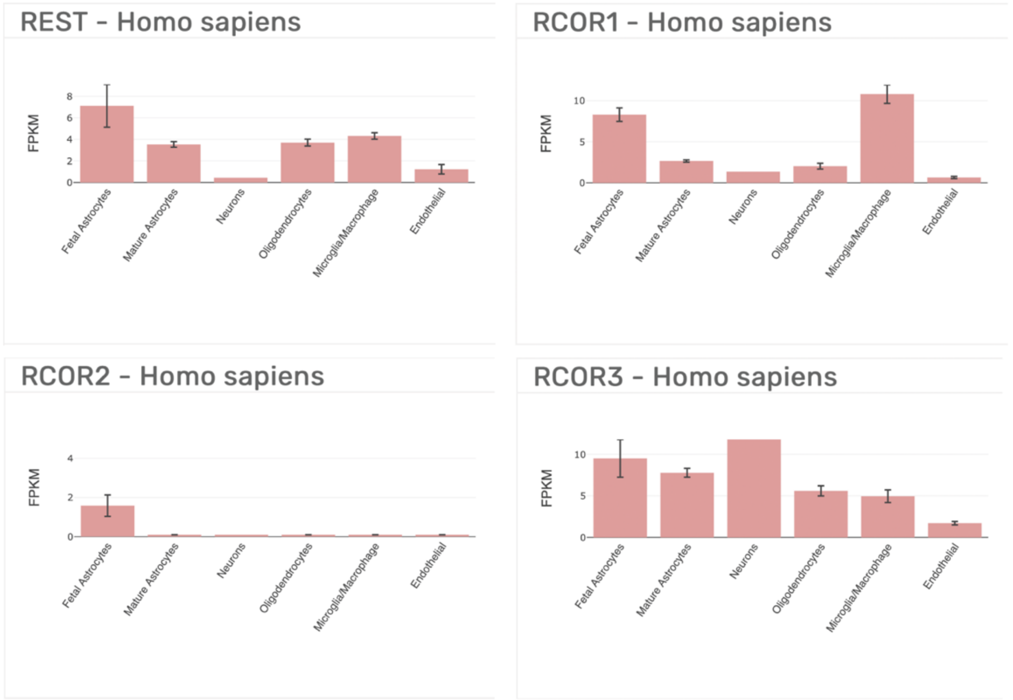

**Figure S7. Cell type specific RNA-seq data of the expression profile of *REST* and *RCOR* genes in human brain.** Data is sourced from the Brain RNA-seq database which is a collection of RNA-seq of purified cell types from human brains [44].

**Table S1. List of genes analysed in Nanostring for neuronal characterisation.**

| **Gene name** | **Approved name** | **Accession** | **Position** | **Category** |
|---|---|---|---|---|
| ***AARS*** | Alanyl-tRNA synthetase | NM_001605.2 | 836-935 | Housekeeper |
| ***ASB7*** | Ankyrin repeat and SOCS box containing 7 | NM_024708.3 | 1281-1380 | Housekeeper |
| ***CCDC127*** | Coiled-coil domain-containing 127 | NM_145265.2 | 295-394 | Housekeeper |
| ***CHAT*** | Choline O-acetyltransferase | NM_020549.4 | 1106-1205 | Cholinergic neuronal marker |
| ***CNOT10*** | CCR4-NOT transcription complex subunit 10 | NM_001256741.1 | 1963-2062 | Housekeeper |
| ***DLX1*** | Distal-less homeobox 1 | NM_001038493.1 | 1336-1435 | Neural progenitor |
| ***DLX2*** | Distal-less homeobox 2 | NM_004405.3 | 591-690 | Neural progenitor |
| ***SLC1A3*** **(EAAT1)** | Solute carrier family 1 member 3 (glial high affinity glutamate transporter 1) | NM_004172.4 | 559-658 | Astrocyte marker |
| ***EID2*** | EP3000-interacting inhibitor of differentiation 2 | NM_153232.3 | 566-665 | Housekeeper |
| ***GAD2*** | Glutamate decarboxylase 2 | NM_000818.2 | 1246-1345 | GABAergic neuronal marker |
| ***GNG3*** | G protein subunit Gamma 3 | NM_012202.1 | 176-275 | Neuronal marker |
| ***GRIA1*** | Glutamate ionotropic receptor AMPA type subunit 1 | NM_000827.3 | 2841-2940 | Neuronal marker |
| ***GRIA2*** | Glutamate ionotropic receptor AMPA type subunit 2 | NM_001083620.1 | 866-965 | Neuronal marker |
| ***GRIN1*** | Glutamate ionotropic receptor NMDA type subunit 1 | NM_000832.5 | 1291-1390 | Neuronal marker |
| ***ISL1*** | Insulin Enhancer protein (ISL) LIM homeobox 1 | NM_002202.2 | 1376-1475 | Neuronal marker |
| ***L1CAM*** | L1 cell adhesion molecule | NM_024003.2 | 3241-3340 | Neuronal marker |

| | | | | |
|---|---|---|---|---|
| ***MAP2*** | Microtubule associated protein 2 | NM_031845.2 | 5171-5270 | Neuronal marker |
| ***MAPT*** | Microtubule associated protein tau | NM_016834.3 | 1206-1305 | Neuronal marker |
| ***MTO1*** | Mitochondrial tRNA translation optimization 1 | NM_133645.2 | 1466-1565 | Housekeeper |
| ***NANOG*** | Nanog homeobox | NM_024865.2 | 1101-1200 | Pluripotency marker |
| ***NDRG4*** | N-Myc downstream regulated 1 | NM_001242835.1 | 3055-3154 | Neuronal marker |
| ***NKX2-1*** | NK2 homeobox 1 | NM_003317.3 | 2012-2111 | Neural progenitor |
| ***PAX6*** | Paired box 6 | NM_000280.3 | 1174-1273 | Neural progenitor |
| ***DLG4*** **(PSD95)** | Disc large MAGUK scaffold protein 4 (post-synaptic density protein 95) | NM_001365.3 | 2461-2560 | Synapse marker |
| ***RABEP2*** | Rabaptin, RAB GTPase-binding effector protein 2 | NM_024816.2 | 1783-1882 | Housekeeper |
| ***RCOR1*** | REST Corepressor 1 | NM_015156.3 | 1726-1825 | CoREST1 |
| ***RCOR2*** | REST Corepressor 2 | NM_173587.3 | 1204-1303 | CoREST2 |
| ***RCOR3*** | REST Corepressor 3 | NM_001136224.2 | 981-1080 | CoREST3 |
| ***REST*** | RE1-Silencing transcription factor | NM_001193508.1 | 1141-1240 | REST |
| ***SUPT7L*** | SPT7 like, STAGA complex gamma subunit | NM_014860.2 | 1171-1270 | Housekeeper |
| ***SYN1*** | Synapsin I | NM_006950.3 | 566-665 | Synapse marker |
| ***TADA2B*** | Transcriptional adaptor 2B | NM_152293.2 | 1589-1688 | Housekeeper |
| ***TH*** | Tyrosine hydroxylase | NM_000360.3 | 1307-1406 | Dopaminergic neuronal marker |
| ***TUBB3*** | Tubulin beta 3 class III | NM_006086.2 | 1538-1637 | Early neuronal marker |
| ***ZNF324B*** | Zinc finger protein 324B | NM_207395.2 | 2821-2920 | Housekeeper |

**Table S2. Cell culture media and components**

| Media | Components |
|---|---|
| Neural induction media (NIM) | 1:1 ratio Neurobasal medium (NBM; Life Technologies, #21103-049) and DMEM/F12 (high glucose) supplemented with 1x N-2 supplement (Life Technologies, #17502-048), 1x B-27 supplement (without vitamin A; Life Technologies, #12587-010), 1x Insulin-Transferrin-Selenium-A (Life Technologies, #51300-044), 2 mM L-Glutamine (Gibco, #25030) and 0.3% glucose (Sigma-Aldrich, #G8769). |

| | |
|---|---|
| Neural media (NM) | NBM supplemented with 1x N-2 supplement 1x B-27 supplement, 1x Insulin-Transferrin-Selenium-A and 2 mM L-glutamine. |
| BrainPhys | Brainphys medium (StemCell Technologies, #05790) supplemented with NeuroCult SM1 (without vitamin A; StemCell Technologies, #05731) and N2 supplement-A (StemCell Technologies, #07152). |
| Embryoid body media (EBM) | DMEM/F12 (ThermoFisher Scientific, #12500-096) supplemented with 1% NEAA (ThermoFisher Scientific, #11140-050), 1% Glutamax (ThermoFisher Scientific, #35050-061), 1% B27 (ThermoFisher Scientific, #17504-001), 1% N2 (ThermoFisher Scientific, #17502-001). |
| Organoid maturation media (OMM) | Brainphys medium containing 1% NEAA, 1% Glutamax, 2% B27, 1% N2. |
| Growth media (GM) | DMEM/F12 media supplemented with 2% B27 supplement (Gibco, #17504044), 1% FBS (Gibco, #10439001), 2 mM Glutamax (Gibco, #35050061), 8 ng/mL FGF2 (Miltenyi Biotec, #130-093-839), 5 ng/mL Ciliary neurotrophic factor (CNTF; Miltenyi Biotec, #130-108-972) and 10 ng/mL Bone morphogenetic protein 4 (BMP4; Miltenyi Biotec, #130-111-165). |
| Maturation media (MM) | DMEM/F12 media supplemented with 1% N2 supplement, 1 mM sodium pyruvate (Gibco, #11360070), 2 mM Glutamax, 5 ug/mL N-acetyl-cysteine (NAC; Sigma-Aldrich, #A9165), 5 ng/mL Heparin-Binding EGF-Like Growth Factor (HB-EGF; Sigma-Aldrich, #E4643), 10 ng/mL CNTF, 10 ng/mL BMP4 and 100 µM DBcAMP (Sigma-Aldrich, #D0627). |

**Table S3. List of primers used for RT-qPCR.**

| Target | Forward primer (5’-3’) | Reverse primer (5’-3’) | Annealing temperature (ºC) | Company |
|---|---|---|---|---|
| ***β2M*** | AAGGACTGGTCTTTCTATCTC | GATCCCACTTAACTATCTTGG | 60 | Sigma-Aldrich |
| ***GAD1*** | CAATACCACTAACCTGCGCC | TTCTCTTCCAGGCTGTTGGT | 60 | Sigma-Aldrich |
| ***GAPDH*** | TCGGAGTCAACGGATTTGGT | TTCCCGTTCTCAGCCTTGAC | 60 | Sigma-Aldrich |
| ***GFAP*** | GCTGGTTTCTCGAATCTG | GAAAAATAGGCCTTGCCTTAG | 58 | KICqStart predesigned SYBR green primers, Sigma-Aldrich |
| ***MAP2*** | CAACGGAGAGCTGACCTCA | CTACAGCCTCAGCAGTGACTA | 58 | Sigma-Aldrich |
| ***PAX6*** | AGAGAATACCAACTCCATCAG | GATAATGGGTTCTCTCAAACTC | 58 | KICqStart predesigned SYBR green primers, Sigma-Aldrich |
| ***PPIA*** | ACGTGGTATAAAGGGGCGG | CTGCAAACAGCTCAAAGGAGAC | 60 | Sigma-Aldrich |
| ***RCOR1*** | CAAGCCATCAGGAAATATGG | CTTCATCTATGTTGAAGCGG | 58 | KICqStart predesigned SYBR green primers, Sigma-Aldrich |
| ***RCOR2*** | CTACTCTTGGAAGAAGACCC | CTCATCACTGTCTTCTTTGTC | 58 | KICqStart predesigned SYBR green primers, Sigma-Aldrich |
| ***RCOR3*** | AGAGGGTAATACTGAACAACC | ATTGGGACTACAGGAAACTG | 58 | KICqStart predesigned SYBR green primers, Sigma-Aldrich |
| ***REST*** | TACTCATTCAGGTGAGAAGC | GTGGGCAATTAAGAGGTTTAG | 58 | KICqStart predesigned SYBR green primers, Sigma-Aldrich |
| ***SLC17A7*** | GAGTTTCGGAAGCTAGCGGG | ACTCAGCTCCAGCGTCTCCG | 60 | Sigma-Aldrich |

| | | | | |
|---|---|---|---|---|
| ***TBR1*** | ACGAACAACAAAGG AGCTTCA | TGGTACTTGTGCAAG GACTGTA | 60 | Sigma-Aldrich |

**Table S4.** Primary antibodies used for immunocytochemistry

| Target | Catalogue number | Dilution |
|---|---|---|
| ALDOC | Abcam, ab190368 | 1:100 |
| CoREST3 | Abcam, ab76921 | 1:200 |
| EAAT2 | Abcam, ab41621 | 1:100 |
| GAD67 | Millipore, MAB5406 | 1:500 |
| GFAP | Sigma-Aldrich, AB5541 | 1:500 |
| MAP2 | Sigma-Aldrich, M4403 | 1:500 |
| NFIB | Abcam, ab186738 | 1:100 |
| S100ß | Abcam, ab52642 | 1:100 |
| SOX9 | Abcam, ab76997 | 1:100 |
| VGLUT1 | Abcam, ab72311 | 1:100 |
